# Soil carbon mineralization and microbial community dynamics in response to PyOM addition

**DOI:** 10.1101/2023.06.21.545992

**Authors:** Nayela Zeba, Timothy D. Berry, Monika S. Fischer, Matthew F. Traxler, Thea Whitman

## Abstract

Wildfires can either negatively impact soil carbon (C) stocks through combustion or increase soil carbon stocks through the production of pyrogenic organic matter (PyOM), which is highly persistent and can affect non-pyrogenic soil organic carbon (SOC) mineralization rates. In this study, we used fine-resolution ^13^CO_2_ flux tracing to investigate PyOM-C mineralization, soil priming effects, and their impacts on soil microbial communities in a Californian mixed conifer forest Xerumbrept soil burned in the 2014 King Fire. We added PyOM produced from pine biomass at 350 °C and 550 °C to the soil and separately traced the mineralization of ^13^C-labeled water-extractable and non-water-extractable PyOM-C fractions in a short-term incubation.

Our results indicate that the water-extractable fraction is 10-50x more mineralizable in both 350 °C and 550 °C PyOM treatments than the SOC or non-water-extractable PyOM fraction. 350 °C PyOM addition led to a short-term positive priming effect, likely due to co-metabolism of easily mineralizable PyOM-C and the SOC, whereas 550 °C PyOM addition induced negative priming, potentially due to physical protection of SOC. We observed significant shifts in bacterial community composition in response to both 350 °C and 550 °C PyOM, with positive PyOM responders belonging to the genera *Noviherbaspirillum*, *Pseudonocardia*, and *Gemmatimonas*. In contrast, fungal communities were less responsive to PyOM additions. Our findings expand our understanding of the post-fire cycling of PyOM and SOC, providing insights into the microbial mineralization of different PyOM-C fractions and their influence on soil C dynamics in fire-affected ecosystems.

## Introduction

Since the mid-1970s, the number of forest wildfires in the western US and the percentage of area burned within them have increased due to a warming climate (1). These fires can impact the biogeochemical cycling of nutrients like carbon (C) within an ecosystem. Increasing fire frequencies can have a negative impact on C storage in forest soils, predominantly due to combustion of wood and litter as well as organic matter in the upper soil layers, leading to direct C losses from the system. Furthermore, the combustion of biomass can also reduce C inputs to soil, which can contribute to the loss of C (2). Simultaneously, fires can have an indirect positive effect on C storage via the production of pyrogenic organic matter (PyOM) (3). For example, during fires in pine-dominated forest systems, between 5-27% of the biomass C can be converted to PyOM, which is likely to persist in soils due to a high proportion of C that is resistant to degradation (3,4). PyOM is also produced intentionally as a soil amendment and as part of a C management strategy, in which case it is referred to as biochar (5). The conditions under which the addition of PyOM/biochar to soils will have a net positive C impact is still not clearly understood and depends on many factors, including the chemical composition and mineralizability of PyOM itself and whether its presence in the soil affects the mineralizability of soil organic carbon (SOC) (6–8). *A key step to addressing this problem is taking a mechanistic approach to understanding how soil microbes decompose different carbon fractions of PyOM in fire-affected soils and if PyOM addition increases the abundance of microbes with the capacity to degrade the different C fractions in PyOM.* This is the overarching goal of this study.

In recent years, through a combination of field studies and lab incubations, we have gained a better understanding of the factors that can affect the mineralization of PyOM by soil microbes. These factors primarily include PyOM properties like feedstock and pyrolysis temperature, PyOM chemical and physical properties, soil characteristics, and the duration of measurement (7,9). These same factors also determine whether PyOM addition will affect the mineralization of the native SOM – *i.e*., whether PyOM addition will induce a “priming” effect (6–8,10). However, a key challenge to making these predictions is that PyOM is not homogeneous, even within a given sample, and different PyOM fractions may play fundamentally different roles in determining its net effect on soil C stocks. For example, a relatively small fraction of PyOM-C, included in the water-extractable fraction, is more easily mineralizable by microbes and may drive the observed increases in SOC mineralization (11–14). Differences in the relative abundance of this easily mineralizable fraction would be reflected in the degree to which PyOM stimulates microbial activity, thereby increasing the rate of SOC mineralization (*i.e.*, inducing a “positive priming effect”) in the short term (6,15). Simultaneously, sorption is often invoked as an explanation for decreased SOC mineralization with PyOM additions (“negative priming”) (6,8), and would be expected to be associated with the more persistent and water-insoluble (*i.e.*, non-water-extractable) PyOM fractions. However, it can be challenging to independently trace the cycling of these different PyOM fractions, as adding them to soil separately would undermine the object of understanding how these fractions cycle upon bulk PyOM addition.

In this study, we aimed to determine if C fractions within PyOM are differentially mineralized by microbes and how this affects the mineralization of SOC. We addressed this question by tracking the mineralization of two ^13^C labeled PyOM-C fractions – ^13^C labeled water-extractable and non-water-extractable PyOM-C in our soil-^13^C PyOM incubations and used this information to decipher the dominant priming mechanisms. We sought to isolate the short-term dynamics of the water-extractable *vs.* non-water-extractable PyOM fraction, given that positive priming effects usually occur over relatively short timescales. We produced PyOM at two different pyrolysis temperatures (350 and 550 °C) to capture the differences in both PyOM-C content and chemistry with increasing temperature, particularly the fraction of water-extractable PyOM-C. PyOM-C concentration increases with higher pyrolysis temperatures, retaining a relatively lower proportion of easily mineralizable PyOM-C structures (16–18). We hypothesized that the water-extractable PyOM-C fraction would be more rapidly mineralized, since it primarily consists of aliphatic C compounds that are readily consumed by microbes, and that the mineralizability of this fraction would be a strong predictor of SOC priming.

While it is important to understand how PyOM chemistry affects its cycling by microbes, it is also critical to understand how PyOM affects the soil microbial communities. There are numerous ways in which PyOM addition can affect soil microbes in the short term, such as the availability of easily mineralizable C as a nutrient source (14), changes in soil properties such as water holding capacity and pH (19,20), and interactions with signaling molecules (21). To date, a relatively small number of lab and field studies have looked at the effects of PyOM on soil bacterial communities (summarized in Woolet and Whitman (22)). While individual studies often found significant effects of PyOM on soil bacterial community composition, soil characteristics played a more important role than PyOM in predicting community composition across soil types (22). In addition to community-wide effects, numerous PyOM-responsive bacterial taxa, across phylogenetic levels have been identified in recent studies. However, only a few genera have been found to have a consistently positive response to PyOM across soil types – these genera include known fire responders, and many have members that are known PAH degraders, indicating a capacity to break down complex aromatic C fractions in PyOM (22). In addition to PyOM-responsive taxa identified via high throughput sequencing, specific bacteria (*e.g.*, *Streptomyces* sp.) readily grow on agar media containing PyOM-C as the sole carbon source (9), while others (*e.g., Pseudomonas* and *Pseudonocardiaceae* sp.) can colonize the PyOM surface (23,24).

Much of our understanding of the effects of PyOM on soil fungi comes from studying fungal responses to fire. Certain fungi, such as *Pholiota*, *Pyronema* and *Penicillium* sp. are known to thrive in post-fire environments (25). Their potential to exploit post-fire resources, such as PyOM, could contribute to their relative increase (26–28), along with other factors like heat tolerance (25,29). A few studies examining the effects of PyOM application on fungi have observed changes in the fungal community structure, primarily driven by alterations in soil properties (30,31). The aromatic C fraction in PyOM may also play an important role in shaping the fungal community composition (32). For instance, dominant post-fire fungus, *Pyronema domesticum* has the capacity to break down complex aromatic C when grown on agar media containing PyOM-C (27). Despite these insights, the effect of PyOM addition on soil fungi, both in terms of whole communities and individual taxa remains relatively unexplored.

While the list of PyOM-responsive fungal and bacterial taxa is growing, the mechanisms by which both the community and individual taxa respond to PyOM are poorly understood. Specifically, with regard to the C heterogeneity of PyOM, it is possible that the relative abundance of some species increases in response to the easily mineralizable PyOM-C fractions, while others increase in response to the more aromatic PyOM-C fractions. Although methods like stable isotope probing would be required to conclusively demonstrate these different responses, this should result in a time-dependent response, with responders to the easily mineralizable PyOM-C fraction increasing early in the incubation period as this fraction is consumed early on, while responders to the aromatic C fractions emerge later. To investigate these dynamics, we performed parallel soil-PyOM incubations and destructively sampled the soils for microbial community profiling at key time points, informed by real-time isotopically partitioned PyOM-C mineralization data. This allowed us to track shifts in microbial community composition and identify PyOM-responsive taxa that increase in relative abundance at key time points during the incubation period. We also used LC-MS analysis to track changes in the soil C chemical profile during these select time points to characterize the nature of carbon compounds available to soil microbes during the incubation. We hypothesized that PyOM addition would result in shifts in both the bacterial and fungal communities within the first few days, tracking the consumption of the easily mineralizable water-extractable fraction. We expected PyOM responders to belong to known fire-responsive and PyOM-responsive genera. We predicted that early responders would mostly be bacteria, while late responders could include both bacteria and fungi that have the ability to degrade complex aromatic C structures, such as *Rhodococcus*, *Sphingomonas* and *Pyronema* sp. (27,33).

## Materials and Methods

### Soil description

To increase the likelihood of finding PyOM-adapted taxa, we targeted a soil with known burn history. Soil was collected in the winter of 2020 from the El Dorado National Forest in California from a region that burned during the 2014 King Fire (38.86953, -120.61322). The soil belongs to the Pilliken series and is a coarse-loamy, mixed, mesic entic Xerumbrept. Topsoil (0-0.1 m) was collected after removing the O-horizon litter layer and stored at -20 °C until it was shipped to Madison, WI on dry ice, after which it was stored at -20 °C until the start of the incubation. We chose to focus on the top 10 cm of mineral soil as this horizon would be expected to have meaningful contact with surface produced PyOM after a wildfire. Standard soil properties (Table 1) were analyzed at the UW Soil and Forage Lab with a sample that was thawed and sieved to < 2 mm.

**Table 1.**
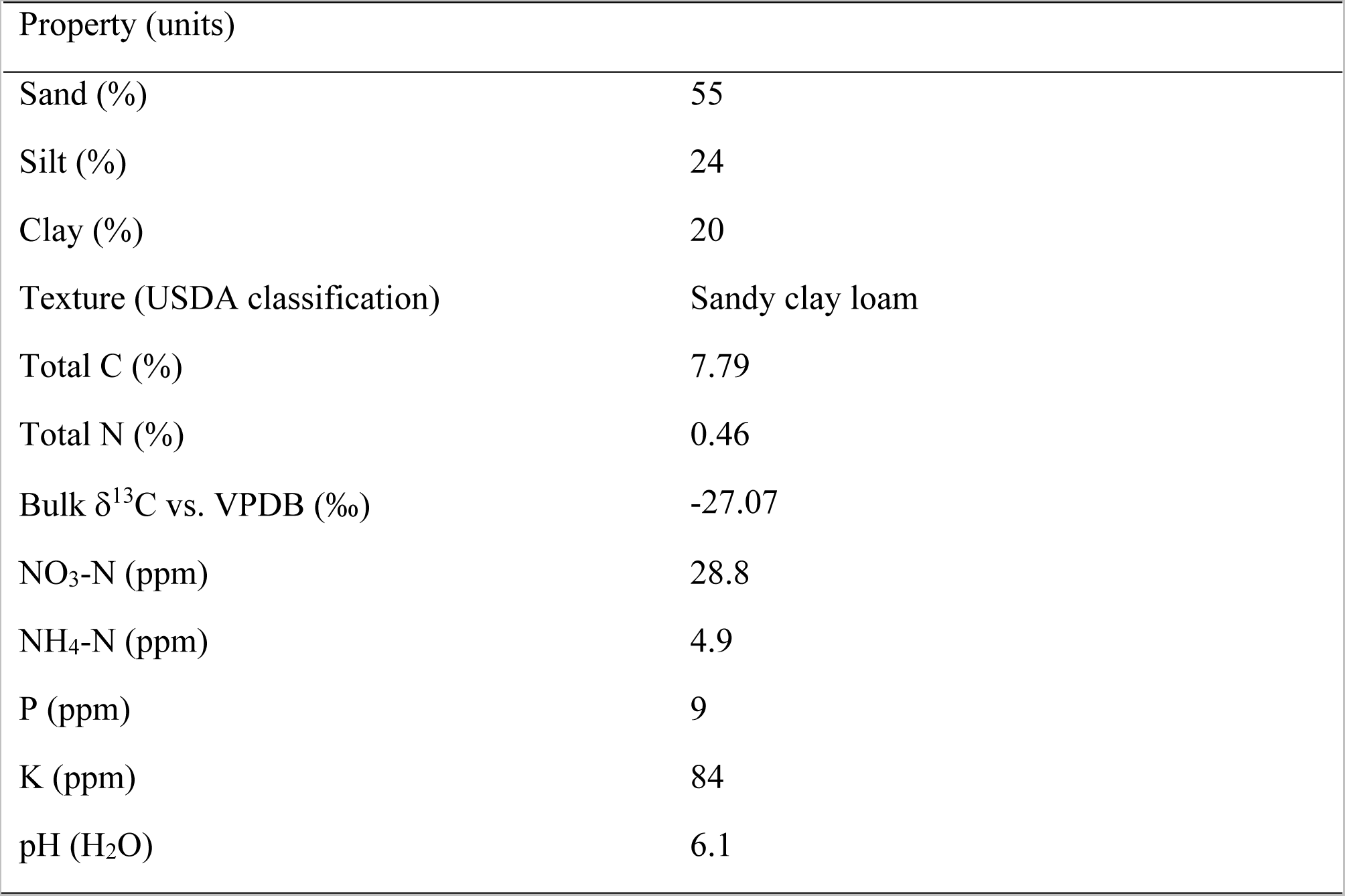
Properties of soil collected from the 2014 King fire affected region

| Property (units) |  |
| --- | --- |
| Sand (%) | 55 |
| Silt (%) | 24 |
| Clay (%) | 20 |
| Texture (USDA classification) | Sandy clay loam |
| Total C (%) | 7.79 |
| Total N (%) | 0.46 |
| Bulk $\delta^{13}\text{C}$ vs. VPDB (‰) | -27.07 |
| NO<sub>3</sub>-N (ppm) | 28.8 |
| NH<sub>4</sub>-N (ppm) | 4.9 |
| P (ppm) | 9 |
| K (ppm) | 84 |
| pH (H<sub>2</sub>O) | 6.1 |

### Biomass production

Two-year-old eastern white pine tree seedlings (*Pinus strobus* (L.)) from the Wisconsin Department of Natural Resources (DNR) were grown in an enriched ^13^CO_2_ atmosphere custom growth chamber for one growing season. The trees were pulse labeled with 99% ^13^CO_2_ at regular intervals over the course of their growth with the goal of producing evenly labeled trees. The labeled trees were watered with deionized water and Hoagland’s solution (Supplemental Note S1).

A paired same-aged set of eastern white pine trees from the Wisconsin DNR, grown under ambient, non-enriched CO_2_, was used as an unlabeled control for this study, keeping moisture, humidity, and light conditions equivalent (Labeled and unlabeled biomass properties are provided in Table S1).

### PyOM production and analyses

We used the aboveground biomass of the eastern white pine trees to produce PyOM. For each set of labeled and unlabeled trees, we ground tree stems and needles and mixed them in 1:4 ratio to account for small differences in the labeled and unlabeled trees. We pyrolyzed both sets of trees at 350 °C (referred to as “350 PyOM”) and 550 °C (referred to as “550 PyOM”) in a modified Fischer Scientific Lindberg/Blue M Moldatherm box furnace (Thermo Fisher Scientific, Waltham, MA, United States) fitted with an Omega CN9600 SERIES Autotune Temperature Controller (Omega Engineering Inc., Norwalk, CT, United States). The PyOM was ground using a mortar and pestle and sieved to collect PyOM with particle size < 45 µm (Additional details in Supplemental Note S2).

Total C, nitrogen (N), bulk d^13^C and d^15^N were measured at the Cornell Stable Isotope Laboratory (COIL) using a Delta V Isotope Ratio Mass Spectrometer (Thermo Fisher Scientific) interfaced to a dry combustion Carlo Erba NC2500 Elemental Analyzer (Carlo Erba Instruments, Milan, Italy). Total hydrogen (H), and oxygen (O) were also measured at COIL on a Delta V Isotope Ratio Mass Spectrometer interfaced to a Temperature Conversion Elemental Analyzer (Thermo Fisher Scientific). The pH was measured in deionized water at a 1:20 solid:solution ratio using an Inlab Micro Combination pH electrode (Mettler Toledo, Columbus, OH, United States) connected to a Thermo Scientific Orion Star A111 benchtop pH meter (Thermo Fisher Scientific). The properties of all PyOM materials are provided in Table 2.

**Table 2.** Properties of PyOM

| Property (units) | $^{13}\text{C}$ labeled | | Unlabeled | |
| --- | --- | --- | --- | --- |
| | 350 $^{\circ}\text{C}$ | 550 $^{\circ}\text{C}$ | 350 $^{\circ}\text{C}$ | 550 $^{\circ}\text{C}$ |
| pH ($\text{H}_2\text{O}$) | 9.42 | 10.09 | 7.97 | 10.19 |
| Total C (%) | $64.75 \pm 2.64$ | $71.87 \pm 1.35$ | $72.87 \pm 8.43$ | $78.17 \pm 1.81$ |
| Total N (%) | $3.24 \pm 0.15$ | $2.91 \pm 0.05$ | $2.82 \pm 0.31$ | $2.47 \pm 0.06$ |
| Total H (%) | 3.52 | 1.75 | 3.78 | 1.93 |
| Total O (%) | 16.28 | 10.96 | 17.46 | 9.32 |
| Bulk $\delta^{13}\text{C}$ vs. VPDB (‰) | $1513.12 \pm 9.47$ | $1596.51 \pm 8.32$ | $-29.38 \pm 0.05$ | $-29.27 \pm 0.08$ |
| Bulk $\delta^{15}\text{N}$ vs. atmospheric air (‰) | $0.40 \pm 0.21$ | $0.46 \pm 0.41$ | $0.50 \pm 0.05$ | $0.74 \pm 0.08$ |
| Total water extractable C (DOC) ($\text{mg g}^{-1}$ PyOM) | 2.11 | 0.95 | 1.24 | 0.67 |

pH values of the original PyOM before adjustment are shown. The values presented for total C, total N, bulk $\delta^{13}\text{C}$ and bulk $\delta^{15}\text{N}$ are means of five replicates ± standard deviation. The values for total water-extractable C represent DOC in original PyOM before exchange.

### Water-extractable PyOM-C extraction and exchange

To isolate and compare the mineralization rates between water-extractable and non-water-extractable PyOM-C fractions, we removed and exchanged the water-extractable fraction from the ^13^C labeled vs. unlabeled PyOM, at C-equivalent rates. This resulted in two PyOM treatments (Fig.1): ^13^C water-extractable PyOM (where the water-extractable fraction is ^13^C-labeled) and ^13^C non-water-extractable PyOM (where the non-water-extractable fraction is ^13^C-labeled). As controls, we extracted and then returned the water-extractable fractions for the ^13^C PyOM and unlabeled PyOM samples at the same rates. After exchange, samples were pH-adjusted to match that of soil and dried before use in the incubation study in order to control for pH effects and isolate C-related phenomena (Additional details in Supplemental Note S3).

**Figure 1.**
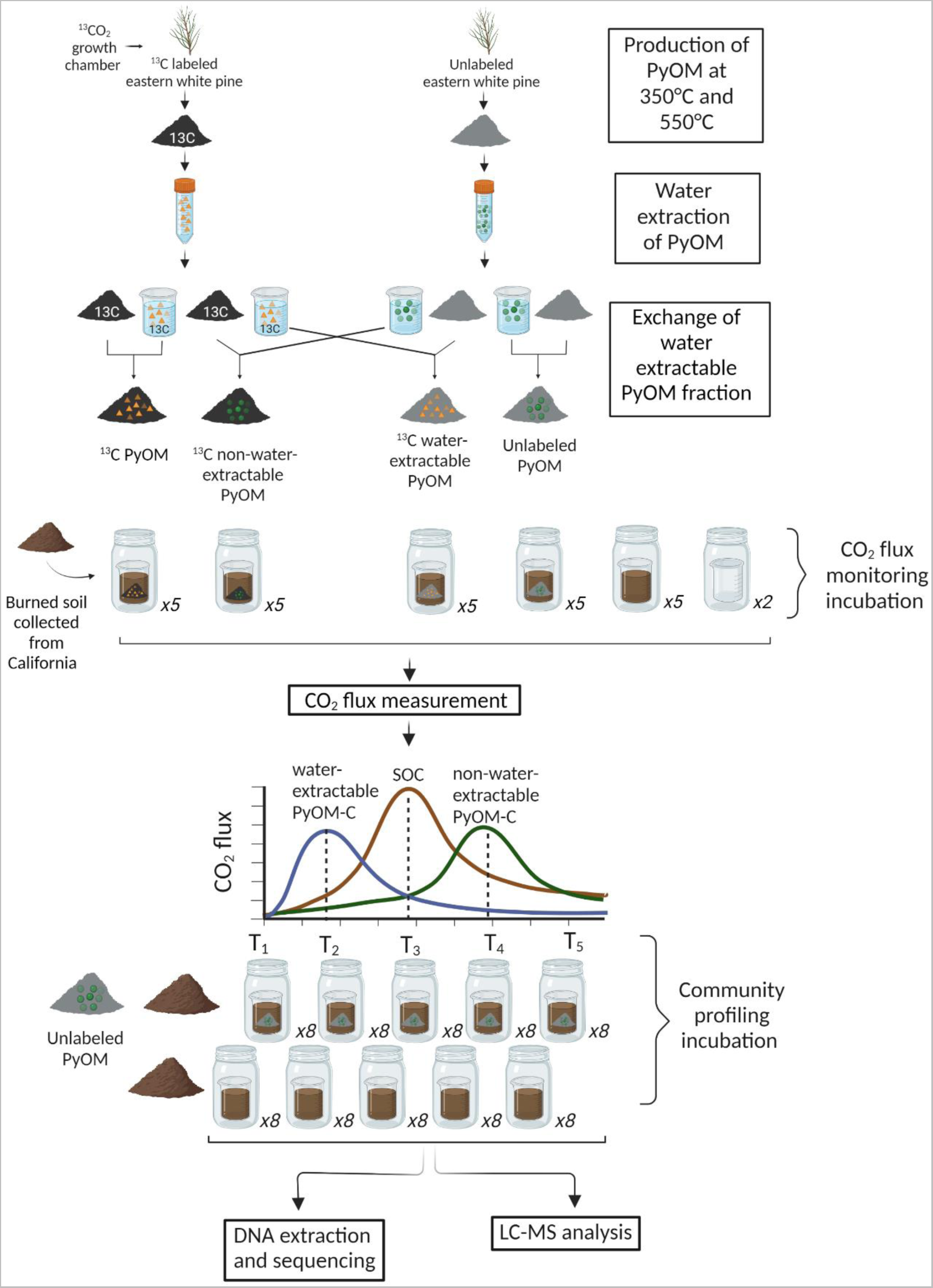
Experimental setup. **A)** Production, extraction and exchange of PyOM. **B)** Incubation set up and monitoring.

Before returning the water-extractable fraction (DOC) to the non-water-extractable PyOM, we dried the non-water-extractable PyOM at 70 °C and measured total C, N, bulk δ^13^C, δ^15^N and pH as described above. We also measured the total organic carbon (TOC) content of the non-water-extractable PyOM, at the UW Soil and Forage Lab using the dry combustion technique (Table S2). We then returned the water-extractable PyOM to the non-water-extractable PyOM fractions (Fig. 1), adjusted the pH of all treatments using HCl additions, and then dried the final materials at 70 °C. The ratios of water-extractable to non-water-extractable PyOM-C were selected to match the DOC content of the water-extracted PyOM for the ^13^C labeled 350 and 550 PyOM treatments (Table 2). Thus, all 350 PyOM treatments had 2.1 mg water-extractable PyOM-C g^-1^ non-water-extractable PyOM-C, while all 550 PyOM treatments had 1 mg water-extractable PyOM-C g^-1^ non-water-extractable PyOM-C.

### Incubation setup and monitoring

Before the incubation, soil was thawed, sieved to < 2 mm and maintained at room temperature and open to the air for two weeks. We used a sub-sample of the soil to determine field capacity for unamended and PyOM amended soils (350 and 550 PyOM) separately to ensure equivalent moisture levels with and without PyOM additions. Water holding capacity was determined as in Whitman et al. (34). The moisture content of the thawed soils was determined one day before the start of the incubation to calculate the water required to reach target moisture levels of 65% field capacity for each treatment.

We ran two sets of parallel incubations – (i) a CO_2_ flux monitoring incubation to partition C mineralization between different PyOM fractions and (ii) a community profiling incubation to analyze the effects of PyOM on microbial community composition at critical time points during the incubation. We analyzed data from the CO_2_ flux monitoring incubation in real time, so we could dynamically select 4-5 timepoints that reflect initial conditions, peak PyOM-C mineralization, peak SOC mineralization and final conditions. At each of these selected timepoints, sample jars from the community profiling incubation were frozen for DNA sequencing. For the flux monitoring jars, we had five replicates for each PyOM temperature and each treatment (Fig. 1): (i) Soil + unlabeled PyOM (ii) Soil + ^13^C PyOM (iii) Soil + ^13^C water-extractable PyOM (iv) Soil + ^13^C non-water-extractable PyOM (v) Unamended soil control. For the community sequencing jars, we were not performing ^13^C partitioning, so we only had two treatments for each temperature, with eight replicates destructively sampled at each timepoint (Fig. 1): (i) Soil + unlabeled PyOM (the same treatment that was used for the flux monitoring incubation) (ii) Unamended soil control.

Incubations were performed in 60 mL glass jars placed inside pint-sized Mason jars (473 mL). Each 60 mL glass jar received 3.5 g soil on a dry mass basis and the soil-PyOM amended jars received PyOM at a consistent rate of 18 mg TOC PyOM g^-1^ dry soil (*i.e.*, 3.1% dry mass addition for 350 PyOM and 2.8% dry mass addition for 550 PyOM). These addition rates were designed to represent locally high inputs of PyOM after a wildfire. We added water dropwise to gradually bring up the moisture of each jar to the target moisture level of 65% field capacity.

### CO2 flux monitoring incubation

After moisture adjustment, we placed the 60 mL glass jars inside Mason jars containing 20 mL acidified deionized water (pH ∼4) at the bottom to maintain humidity and prevent water loss. We capped and sealed the jars with sterile, gas-tight lids with fittings for CO_2_ gas measurements and connected them to randomly selected positions on the distribution manifolds (multiplexer) using polyurethane tubing (35). The connected jars were immediately flushed with a 400 ppm CO_2_-air gas mixture to reset the headspace CO_2_ concentration in all jars at the initial timepoint. We incubated the jars at room temperature in the dark and measured the concentration of CO_2_ emitted in the headspace of each jar at frequent intervals using a Picarro G2131i cavity ringdown spectrometer (Picarro Inc., Santa Clara, CA, United States) attached to the multiplexer over a period of one month. For 350 PyOM, we measured headspace CO_2_ concentration at intervals of 6 h during the first 2 days, and gradually increased the intervals to 12 h during the first and second weeks, 24 h during the third week and 48 h during the last week of incubation. Similarly, for 550 PyOM, we measured headspace CO_2_ concentration every 6-12 h during the first 3 days, 24 h during the first and second weeks and 48-72 h till the end of the incubation. After each measurement, we flushed the jars with the 400 ppm CO_2_-air gas mixture to prevent oxygen depletion and excessive CO_2_ accumulation inside the jars. The precise concentration after flushing each jar was measured and subtracted from the next timepoint reading to calculate the emitted CO_2_ in the jar during that interval.

### Community profiling incubation

For community profiling, after moisture adjustment, we placed the glass jars inside Mason jars containing 20 mL deionized water at the bottom and capped them with sterile regular lids. The jars were then incubated in the dark at room temperature. During the incubation, the jars were opened for 1-2 min every 48 hours to prevent oxygen depletion inside the Mason jars and to mirror the CO_2_ flux incubations. We checked the mass of the jars once a week to determine moisture loss and water was added to return jars to target moisture levels. At each sampling timepoint, eight jars were destructively sampled and placed in sterile Whirl-Pak bags (Nasco, Fort Atkinson, WI, United States) and stored at -80 °C until DNA extraction was performed. During the setup, eight unamended and PyOM amended soil samples were randomly selected and frozen to represent community profile on Day 0.

### ^13^CO_2_ partitioning and statistical analyses

We used R Studio (v4.2.2) (36) with the ‘tidyverse’ (37), ‘zoo’ (38) and ‘broom’ (39) packages to process raw CO_2_ readings from the multiplexer-Picarro system. Stable isotope partitioning, as represented by Equation 1, was used to partition the CO_2_ emissions from the flux monitoring treatments to determine the fraction of CO_2_ emitted from water-extractable PyOM, non-water-extractable PyOM, and soil in the soil-PyOM amended incubations.

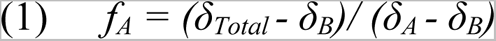

Equation 1 calculates the fraction of CO_2_ emitted from source A (*f_A_*) using the ^13^C isotopic composition of the total respired CO_2_ (*δ_Total_*), CO_2_ respired from source A (*δ_A_*), and source B (*δ_B_*).

For example, we partitioned the total CO_2_ emissions from the “Soil + ^13^C water-extractable PyOM” treatments into two sources: CO_2_ emitted from the ^13^C labeled water-extractable PyOM source and CO_2_ emitted from the unlabeled soil and non-water-extractable PyOM source. We calculated the fraction of total CO_2_ emitted from the water-extractable PyOM source (*f_water-extractable PyOM_*) using Equation 2:

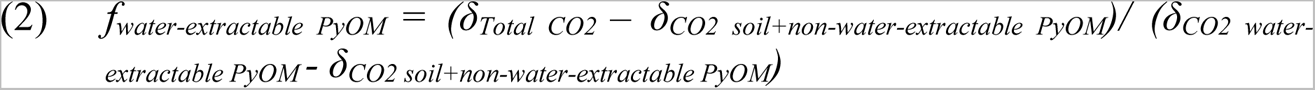

In this equation,

*δ_Total CO2_* represents the mean isotopic composition of the total CO_2_ emitted from the "Soil + ^13^C water-extractable PyOM" treatments, *δ_CO2 soil+non-water-extractable PyOM_* represents the mean isotopic composition of the CO_2_ emitted from soil and non-water-extractable PyOM (determined from the “Soil + unlabeled PyOM” treatments), and *δ_CO2 water-extractable PyOM_* represents the isotopic composition of the CO_2_ emitted from water-extractable PyOM, assumed to be the isotopic composition of the ^13^C labeled PyOM.

Similarly, we determined the fraction of CO_2_ emitted from non-water-extractable PyOM (*f_non-water-extractable PyOM_*) and the fraction of CO_2_ emitted from SOC (*f_SOC_*) by partitioning the flux from other treatments (Supplemental Note S4). We note that the isotopic composition of the labeled water-extractable and non-water-extractable PyOM-C fractions may differ slightly; however, small differences in isotopic composition of the bulk ^13^C-labeled PyOM for CO_2_ emitted from both fractions would not meaningfully impact our overall partitioning results (Fig. S1).

We determined the mineralizability of each PyOM-C fraction by normalizing the amount of C mineralized with the quantity of each carbon source added to the jars. We assessed normality and homogeneity of variance across treatment groups using the Shapiro and Bartlett test functions in the R ’stats’ package. Welch’s ANOVA was employed to test significant differences in mineralizability between the PyOM-C fractions and SOC, due to unequal variances. The Games-Howell post-hoc test function in the ’rstatix’ package (40) was used to compare mineralizability between different treatment pairs. All code used for flux partitioning, data analyses and figures in this manuscript is available at https://github.com/nayelazeba/Chapter3.

### Microbial community analyses

#### DNA extraction, amplification, and sequencing

We extracted DNA from each soil sample using the DNEasy PowerLyzer PowerSoil DNA extraction kit (QIAGEN, Hilden, Germany), following the manufacturer’s instructions. One blank extraction without soil was included for every 24 samples. We performed PCR in triplicate to amplify the extracted DNA, targeting the 16S rRNA gene v4 region with 515f and 806r primers (41), and targeting the ITS2 gene region with 5.8S-Fun and ITS4-Fun primers (42) with barcodes and Illumina sequencing adapters added as per Kozich et al. (43). During PCR, we included one negative control (PCR-grade water) and one positive control (known microbial community mix) for every 30 samples. The PCR amplicon triplicates were pooled, purified and normalized using a SequalPrep Normalization Plate (96) Kit (Thermo Fisher Scientific). Samples, including blanks, were pooled and library cleanup was performed using a Wizard SV Gel and PCR Clean-Up System A9282 (Promega Corporation, Madison, United States) according to manufacturer’s instructions. A detailed procedure for the DNA extraction and PCR amplification and purification is described in Whitman et al. (28). We submitted the pooled library to the UW Madison Biotechnology Center for 2x300 paired end Illumina MiSeq sequencing (Illumina Inc., San Diego, CA, United States) for both 16S and ITS2 amplicons.

#### Sequence data processing and taxonomic assignments

We used the QIIME2 pipeline (QIIME2, v2020.6 (44)) to process the sequences following the steps described in Woolet and Whitman (22). The sequence processing steps were performed on the UW-Madison Centre for High Throughput Computing cluster. Raw sequence data were demultiplexed and quality filtered followed by denoising with DADA2 (45). The dada2 denoise-paired command as implemented within QIIME2 was used to determine operational taxonomic units (OTUs). This resulted in a retention of a mean of 45,246 16S sequences and a mean of 27,358 ITS2 sequences per sample. For ITS2 reads, we then ran the sequences through ITSx (46) to identify fungi and to remove plant and other eukaryotic sequences. Taxonomy was assigned to the 16S sequences using the Silva 138 database (47) at 99% similarity using the QIIME2 feature-classifier classify-sklearn. Pre-trained taxonomy classifiers specific to the primers used for 16S sequencing were used (48). For the IT2 reads, we assigned taxonomy using the UNITE “developer” database (v8.3) at 99% similarity (49). UNITE taxonomy classifiers were trained on the full reference sequences using the QIIME2 feature-classifier fit-classifier-naive-bayes (50).

#### Statistical analyses

We worked in R Studio, primarily with the ‘phyloseq’ (51) and ‘vegan’ packages (52), to compare microbial community composition between soil and PyOM amended soil samples. We removed 381 OTUs that belonged to “Chloroplast” and “Mitochondria” in the 16S OTU table and removed samples with less than 6,650 16S reads and 1,560 ITS2 reads. We then calculated Bray-Curtis dissimilarity between samples (53), normalized by relative abundance and represented them as NMDS ordinations. We tested for significant effects of PyOM addition after controlling for time using permutational multivariate ANOVA (PERMANOVA) on Bray-Curtis dissimilarities, implemented in vegan as the ‘adonis2’ function.

To identify PyOM-responsive taxa, we calculated log_2_-fold changes in taxon abundances in control vs. PyOM-amended soils using the ‘DESeq2’ R package which is used to analyze differential abundance between treatments (54). To test for specific effects of PyOM addition on taxon abundance, we used a design formula that models differences in taxon abundance across samples on Day 0, over time and due to PyOM addition over time (55). We then performed a likelihood ratio test with a reduced model without the PyOM addition effects over time to identify significant responder OTUs (Benjamini and Hochberg correction, adjusted p value < 0.05). This approach allowed us to identify only those taxa that at one or more time points after Day 0 showed a significant log_2_-fold change with PyOM addition.

The t-test function from the ‘stats’ R package was used to test for significant differences in relative abundances between PyOM-amended and unamended soils for PyOM-responsive genera identified using DESeq2.

### LC-MS analysis

To acquire chemical profiles, we first prepared chemical extracts by combining 0.4 g of soil with 4 mL of LC/MS-grade methanol, sonicated for five minutes, and then shaken overnight (∼16 hours) at 25 °C and 200 rpm. Blank extraction controls were prepared in parallel, in which empty tubes lacking any soil sample were subjected to the same chemical extraction protocol. Solids were allowed to settle to the bottom for 30 minutes and then 3.5 mL was carefully collected from the top and immediately dried via Savant SPD1010 SpeedVac Concentrator (Thermo Fisher Scientific). To analyze these dried extracts via LC/MS, we resuspended them in 1 mL of 100 nM nonactin LC/MS-grade methanol solution, to a final concentration of approximately 1 mg extract / 1 mL of solvent. These resuspended samples were sonicated for 5 min to ensure that the extract dissolved into the solvent, and then centrifuged at 15,000 rpm to pellet any particulate, after which, 900 μL of solution was transferred to an HPLC vial. To create a pooled quality control (QC) sample we combined 10 μL of each sample. Samples were analyzed in a randomized order with a methanol blank and pooled QC analyzed after every 12 samples. Samples were analyzed with an ultra-high-pressure liquid chromatography (UHPLC) system Dionex Ultimate 3000 (Thermo Fisher Scientific) coupled to a high-resolution mass spectrometer (HRMS) Thermo Q-Exactive Quadrupole-Orbitrap (Thermo Fisher Scientific) using a heated electrospray ionization (HESI) source and a C18 column (Thermo Scientific Acclaim rapid-separation liquid chromatography [RSLC] system, 50 mm by 2.1 mm, 2.2 μm pore size). We used the following 20 min UHPLC method; 1 min 40% acetonitrile (ACN) plus 0.1% formic acid (FA), 1 min gradient from 40% to 65% ACN plus 0.1% FA, 11 min gradient from 65% to 99% ACN plus 0.1% FA, 3.5 min 99% ACN plus 0.1% FA, 0.5 min gradient from 99% to 40% ACN plus 0.1% FA, and 3 min re-equilibration in 40% ACN plus 0.1% FA; injection volume of 5 μl, flow rate of 0.4 mL/min, and column oven temperature of 35 °C. The full MS1 scan was performed in positive mode at a resolution of 35,000 FWHM (full width at half-maximum) with an automatic gain control (AGC) target of 1e6 ions and a maximum ion injection time (IT) of 100 ms with a mass range from m/z 200 to m/z 2000. Data were processed using MS-DIAL v4.9 (56). We used R v4.1.3 to omit features with a peak height value greater than 100,000 in any negative control samples (*i.e.*, methanol blanks and blank extraction controls) prior to ordination and statistical analyses as described above.

## Results

### PyOM-C and SOC flux dynamics

Over the 30-day incubation, the mineralizability of PyOM-C from the water-extractable fraction was significantly higher than that of the non-water-extractable PyOM-C fraction and soil organic carbon for both the 350 and 550 PyOM treatments (p < 0.05, Games-Howell post-hoc test; Fig. 2.A). The amount of C mineralized per gram of initial C from the water-extractable fraction was 10 and 50 times higher than the non-water-extractable PyOM-C in the 350 and 550 PyOM treatments, respectively. The C mineralizability of the 350 non-water-extractable PyOM-C fraction was higher than that of soil organic carbon. However, this pattern was not observed in the 550 PyOM treatment, where the C mineralizability of soil organic carbon was higher. Our focus in this study was to compare the relative mineralizability of the different PyOM-C fractions with soil organic carbon (*i*.*e*., on a per carbon gram basis). Considering the total C mineralized, the cumulative mineralization from the water-extractable PyOM-C fractions was lower compared to the non-water-extractable PyOM-C and SOC due to the small fraction of total PyOM represented by water-extractable PyOM-C in the jars (Fig. S2).

**Figure 2.**
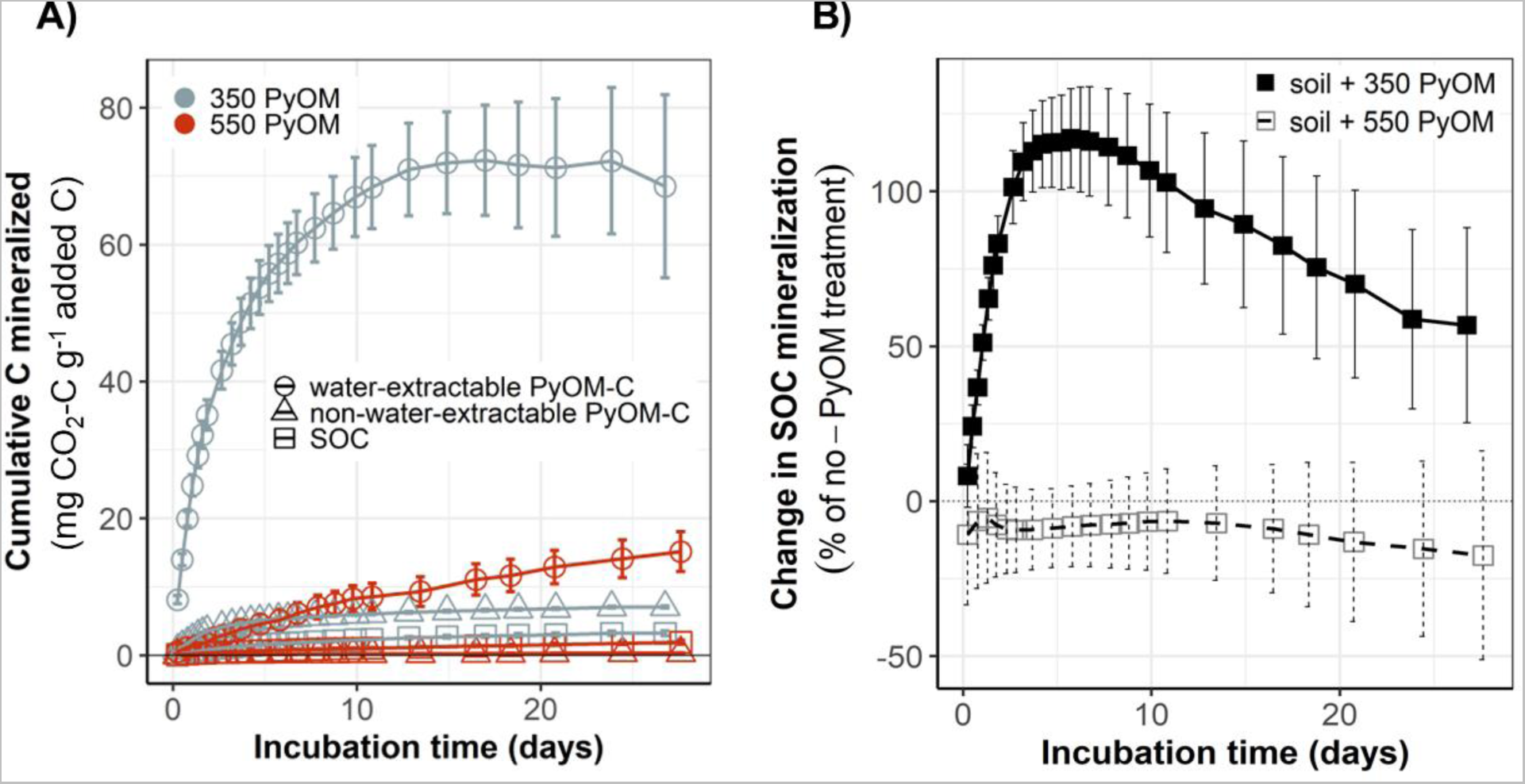
A) Mineralizability of PyOM fractions and SOC over time. Cumulative mean C mineralized per gram of added C for water-extractable PyOM-C, non-water-extractable PyOM-C and SOC. (n=4-5, error bars=SE). **B) Effect of PyOM addition on SOC mineralizability.** Cumulative mean SOC priming over time represented as % of mineralization in no-PyOM treatment. Priming calculated as: [[SOC_(PyOM)_-SOC_(no-PyOM)_] / SOC_(no-PyOM)_] (n=4-5, error bars=SE).

The addition of 350 PyOM resulted in a net 57% increase in cumulative SOC mineralization after 30 days of incubation, indicating a strong positive priming effect (Fig. 2.B). This effect was immediately apparent, with cumulative SOC mineralization in the 350 PyOM amended soils being over 100% higher than in the unamended soils after just 5 days of incubation. However, this strong positive priming effect diminished after 5 days, although the net effect remained positive for the duration of the study. In contrast, the addition of 550 PyOM resulted in a net 17% decrease in cumulative SOC mineralization, indicating a negative priming effect. This negative priming was also observed immediately upon the start of the incubation, with the cumulative amount of SOC mineralized in the 550 PyOM amended soils being on average 8% lower than in the unamended control soil treatments during the first 10 days of incubation.

### Microbial community composition

We observed significant shifts in the bacterial and archaeal community composition over time (PERMANOVA, *p_time_* < 0.01) and in response to the addition of 350 PyOM (PERMANOVA, *p_350PyOM_* = 0.01, R^2^_350PyOM_ = 0.03, Fig. 3.A) and 550 PyOM (PERMANOVA, *p_550PyOM_* < 0.01, R^2^_550PyOM_ = 0.04, Fig. 3.C). The impact of PyOM was evident within a few days of incubation – the communities in both the 350 and 550 PyOM amended soils were distinct from the unamended soils by Day 2 and Day 4, respectively. In contrast, we only detected significant shifts in the fungal community composition over time (PERMANOVA, *p_time_* < 0.01) and not in response to PyOM addition (Fig. 3.B and 3.D). Time had the most explanatory power for variations in both the bacterial and fungal communities (R^2^ values between 0.08 and 0.25).

**Figure 3.**
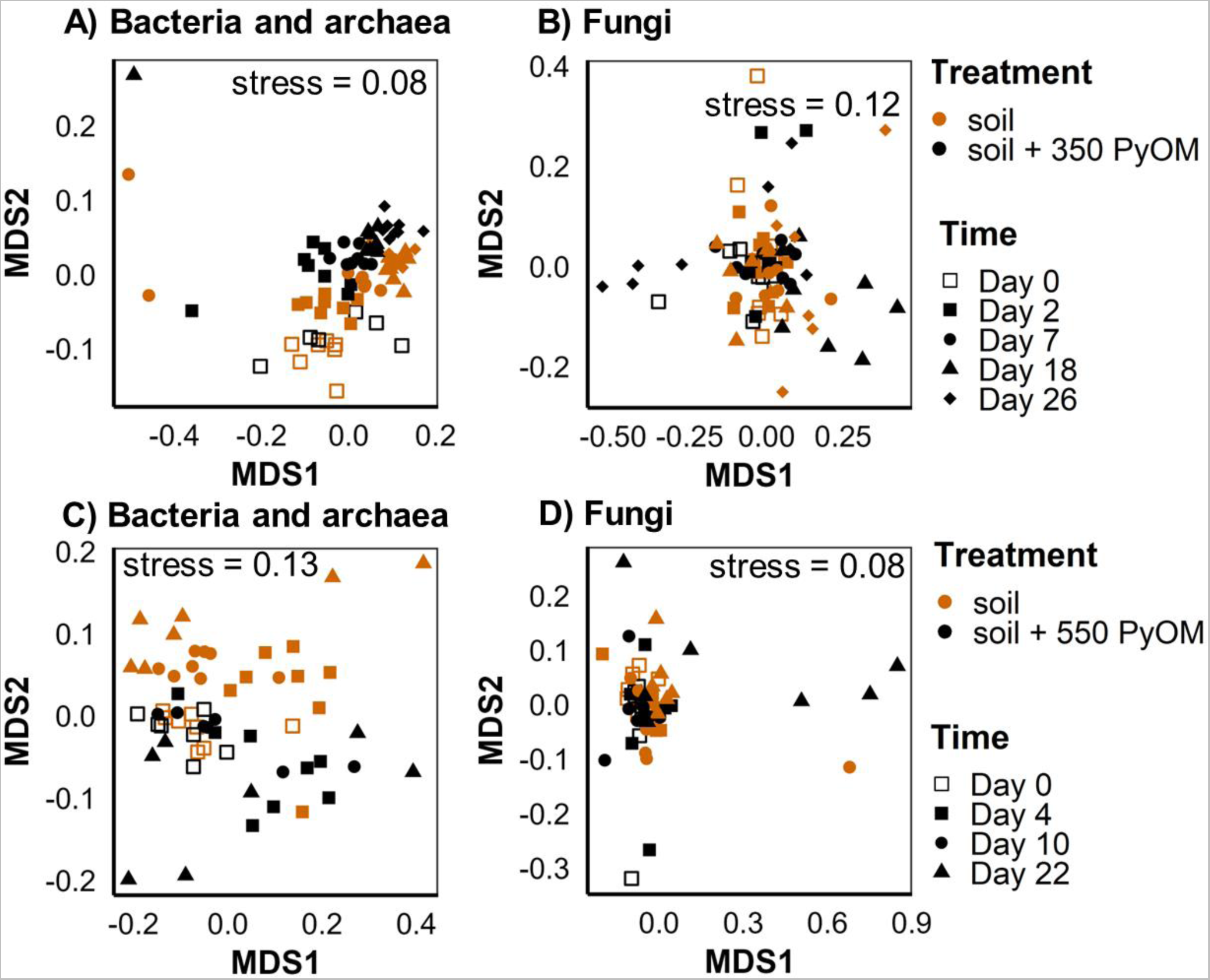
Effect of PyOM addition on soil microbial community composition. NMDS ordination of Bray-Curtis dissimilarities between **(A & C)** bacterial/archaeal (16S rRNA gene v4 region) communities (k=2) and **(B & D)** fungal (ITS2) communities (k=3) for all samples. Top panels indicate data for unamended and 350 °C PyOM-amended soil samples. Bottom panels indicate data for unamended and 550 °C PyOM-amended soil samples.

### Soil C profile

The LC-MS analysis of PyOM amended and unamended soils showed that the addition of both 350 and 550 PyOM resulted in significant shifts in soil C profile (PERMANOVA, *p_350PyOM_* < 0.01, R^2^_350PyOM_ = 0.05; *p_550PyOM_* R^2^_550PyOM_ = 0.06; Fig. 4.A and 4.B).

**Figure 4.**
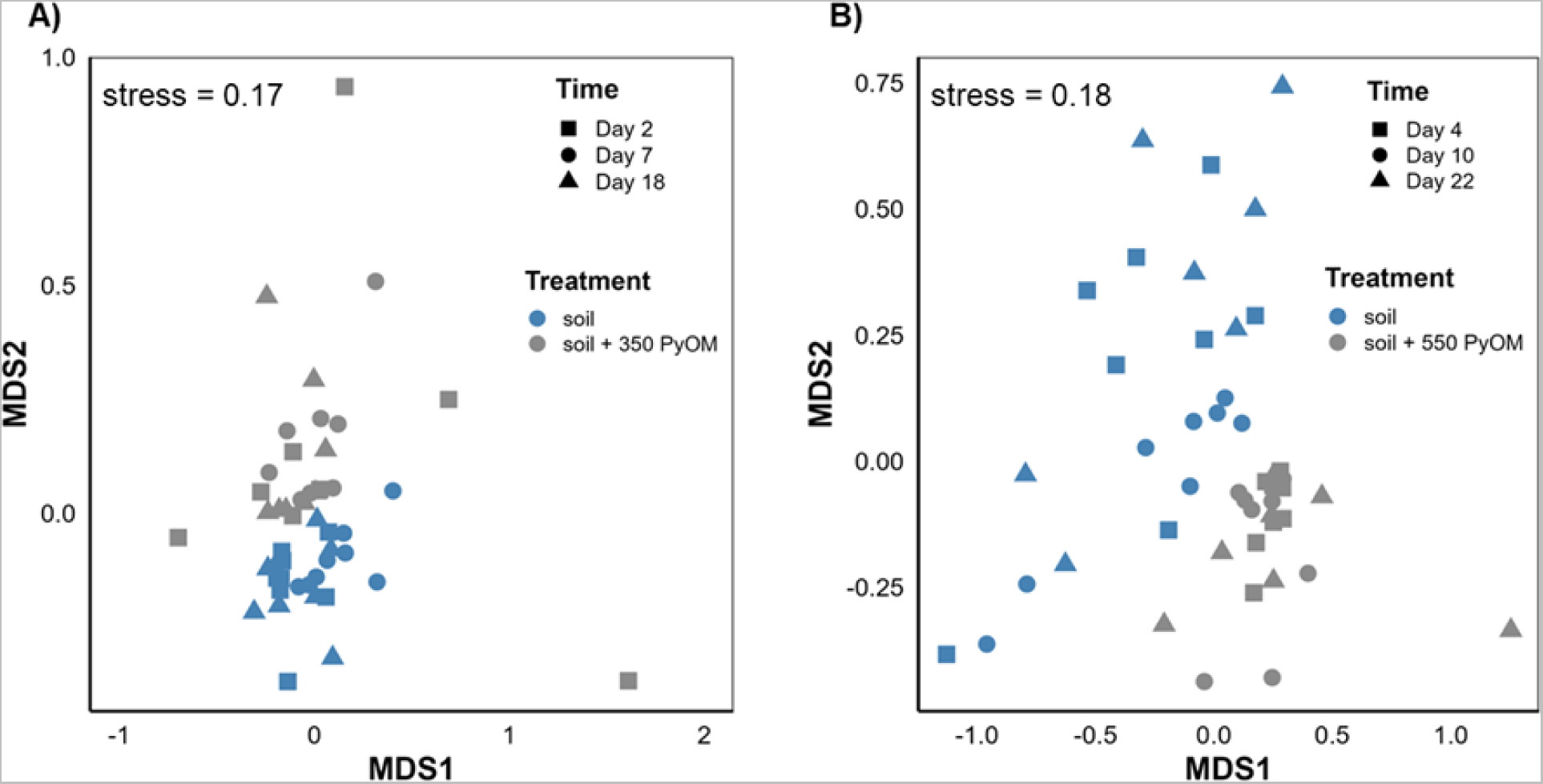
Effect of PyOM addition on soil carbon chemical profile. NMDS ordination of Bray-Curtis distances between soil chemical peaks for **A)** unamended and 350 °C PyOM amended soil samples (k=2) and **B)** unamended and 550 °C PyOM amended soil samples (k=2).

### PyOM positive responders

Using DESeq2, we identified 19 bacterial OTUs that responded positively to the addition of 350 PyOM (Fig. 5.A). These OTUs showed a significant positive response to 350 PyOM after controlling for time and sample effects and had a mean normalized count higher than the 25^th^ percentile. Among these responsive OTUs, 12 belonged to the following genera: *Bacillus*, *Massilia*, *Ferruginibacter*, *Gemmatimonas*, *Noviherbaspirillum*, *Pseudonocardia*, *Psychroglaciecola*, *Saccharimonadales*, and *Singulisphaera*. The remaining 7 OTUs were assigned to unknown genera. For *Gemmatimonas* and *Noviherbaspirillum,* we observed a significant increase in the relative abundance of all responsive OTUs following the addition of 350 PyOM, with the increase appearing at different points during the incubation period (Fig. 5.B). In the case of fungi, we identified 5 OTUs that showed a significant positive response to 350 PyOM and had a mean normalized count higher than the 25th percentile (Fig. S3). These OTUs belonged to the following genera: *Calyptrozyma*, *Coniochaeta*, *Holtermanniella*, *Leucosporidium* and *Solicoccozyma*. However, we did not observe an increase in relative abundance over time for any of the responsive fungal OTUs following the addition of 350 PyOM. Upon 550 PyOM addition, we identified only a single bacterial OTU from the *Gemmatimonas* genus and a single fungal OTU from the *Paraphoma* genus that responded positively to its addition.

**Figure 5.**
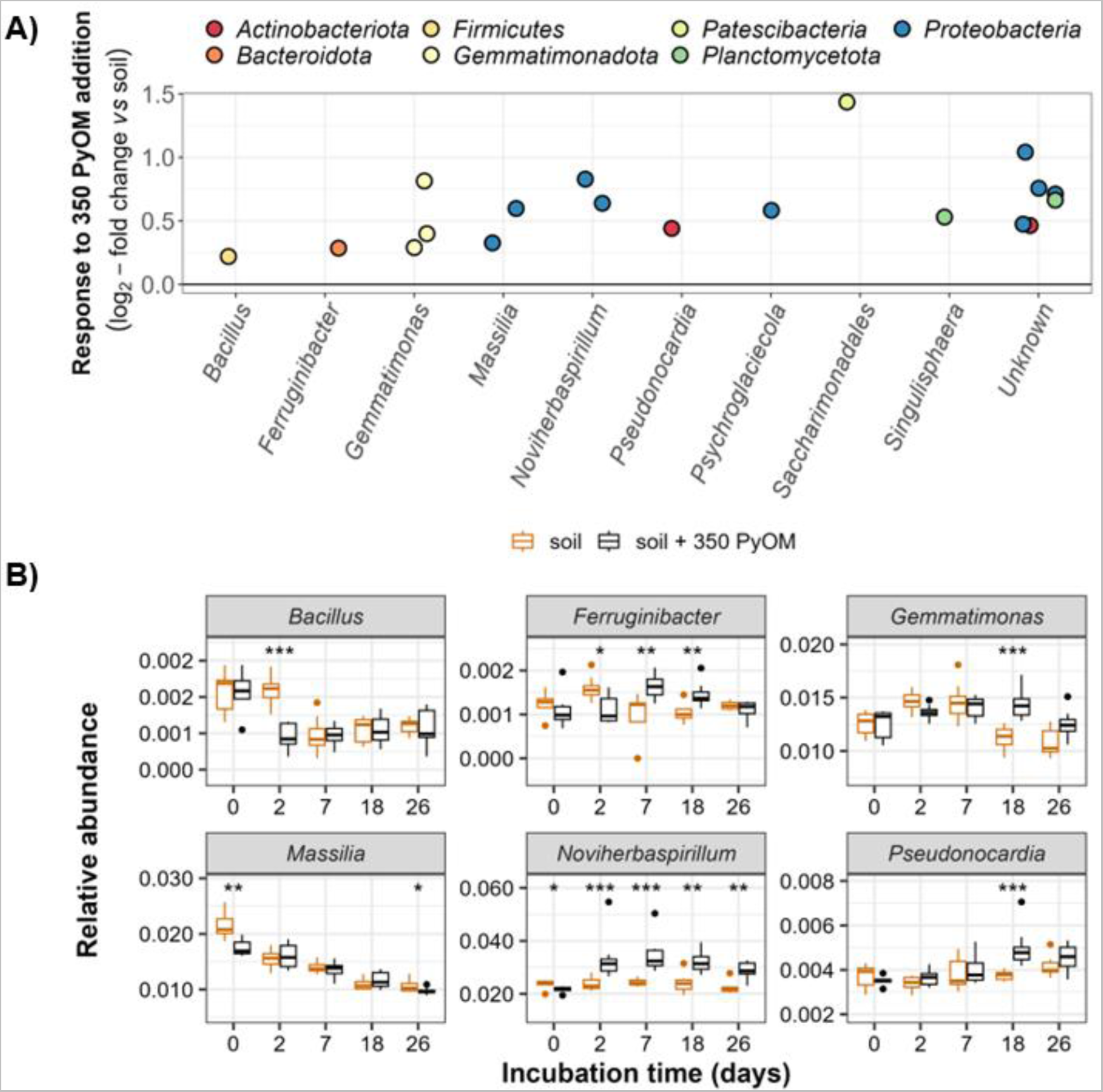
Bacterial response to 350 PyOM. **A)** Log2-fold change in 350 PyOM amended vs. unamended soils, controlling for differences in taxon abundance across samples on Day 0 and over time. Each point represents a single 16S rRNA gene v4 region OTU with mean normalized count above the 25^th^ percentile and that was significantly different in abundance in PyOM amended vs. unamended soils (Benjamini and Hochberg correction, adjusted p value < 0.05). **B)** Relative abundance of six positive responsive genera over time (as identified using DESeq2) observed in the unamended and 350 PyOM-amended soils (n=5-8). Data are grouped for multiple responder OTUs within a genus. * indicates relative abundances that differ significantly from unamended soil at a given timepoint (t-test, *: p < 0.05; **: p < 0.01; ***: p < 0.001).

## Discussion

### Mineralizability of PyOM differed between C fractions and with PyOM temperature

Consistent with our hypothesis, water-extractable C fractions in PyOM were much more readily mineralized than non-water-extractable C fractions (Fig. 2.A). This 10-50x difference in C mineralizability underscores the heterogeneity of the C in PyOM and the potential to explain significant variation in PyOM decomposition rates. Meaningful differences also exist between equivalent fractions of PyOM produced at different temperatures: when we compare the C mineralization between the two different temperatures of PyOM, we see that the mineralizability of both the 350 water-extractable and non-water-extractable fractions was consistently higher. This indicates that C in low-temperature PyOM is more readily decomposed by microbes, possibly due to its lower aromaticity and lower degree of condensation compared to the C in the higher temperature 550 PyOM (17,57) in addition to having a larger fraction of water-extractable C. Interestingly, compared to SOC, microbes preferred the water-extractable PyOM-C in both 350 and 550 PyOM. However, when it came to the non-water-extractable PyOM-C fractions, the non-water-extractable PyOM-C in 350 PyOM was preferred over SOC, while SOC was preferred over 550 PyOM-C. This further highlights the typically higher degrees of condensation in 550 PyOM, making the C more resistant to microbial breakdown.

### Mechanisms of positive and negative priming differed over time and with PyOM temperature

The addition of 350 PyOM caused a positive priming effect on the mineralization of SOC, while the addition of 550 PyOM resulted in a negative priming effect (Fig. 2.B). The positive priming effect of 350 PyOM is likely due to co-metabolism / increased microbial activity, where the addition of easily mineralizable PyOM-C increases total microbial activity and accelerates the mineralization of SOC over short periods of time (6,10,14). This is strongly supported by the significant positive correlation (R^2^ = 0.97, *p* < 0.001) between the rate of PyOM-C mineralization and the rate of SOC priming observed during the incubation (Fig. 6.B). However, the high-frequency sampling afforded by our multiplexer cavity ringdown spectroscopy (CRDS) system allowed us to also detect a negative correlation between these two variables in the first 48 hours (Fig. 6.A). We propose that this is likely due to substrate switching. Substrate switching occurs when microbes preferentially use the easily mineralizable PyOM-C over SOC and can explain negative priming effects in the early stages of incubation (10,15). The higher C mineralizability observed from both fractions of 350 PyOM compared to SOC, particularly in the first 48 hours, supports the argument that the added PyOM-C is a more favorable substrate than the existing SOC, largely driven by the most available constituents of the water-extractable fraction in 350 PyOM. This preferential usage within the first two days results in a scenario where the remaining carbon in 350 PyOM and SOC are both readily used by microbes through the remaining incubation period, resulting in a net positive priming effect. Positive priming could also be a result of increased microbial activity due to alleviation of nutrient constraints or soil conditioning (creation of favorable microenvironments) upon PyOM addition (10,58).

**Figure 6.**
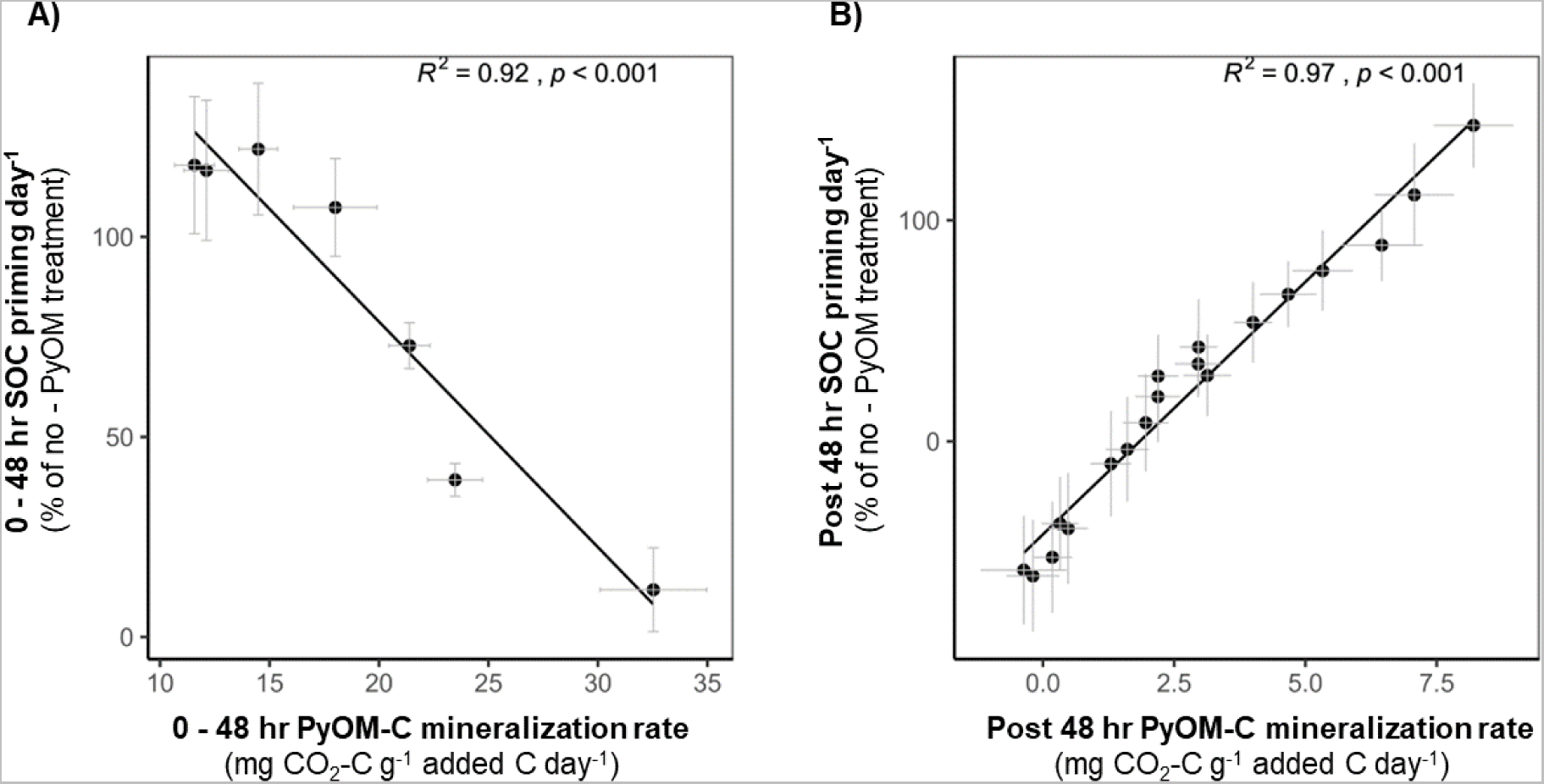
Relationships between mean PyOM-C mineralization rate and SOC priming for 350 °C PyOM. **A)** 0 - 48 hours of soil-PyOM incubation and **B)** post 48 hours of soil-PyOM incubation. (n=4-5, error bars=SE). Priming calculated as: [[SOC_(PyOM)_-SOC_(no-PyOM)_] / SOC_(no-PyOM)_].

We can rule out substrate switching as an explanation for the negative priming effect of 550 PyOM on SOC mineralization, as no negative correlation was observed between the rate of 550 PyOM-C mineralization and the rate of SOC priming (Fig. S4). Instead, the short-term negative priming effect may be due to inhibition, sorption of SOC on PyOM, or dilution. Inhibitory effects of 550 PyOM on microbes (such as reduction in microbial biomass) were not investigated, but cannot be ruled out as a potential cause. Inhibition is known to occur indirectly through changes in the soil environment or directly due to toxic chemicals released upon the addition of PyOM that inhibit microbial activity (59,60). However, studies investigating the impact of PyOM produced at varying pyrolysis temperatures on microbial populations have not found a significant reduction in microbial biomass with increasing pyrolysis temperature (10,61), making inhibition an unlikely explanation in this study. Sorption of SOC on high-porosity, high-surface-area 550 PyOM, may also contribute to the negative priming effect by making SOC less accessible to microbes (10). Dilution of the SOC pool by the addition of PyOM-C, even with just a small mass of easily mineralizable C, may also decrease mineralization. In a previous study, DeCiucies et al. (10) found that dilution contributed to 19% of reductions in SOC mineralization observed with PyOM produced at 450 °C over the first 7 days. The high PyOM addition rate (compared to the range of additon rates in Wang et al. (7) and low C content in the water-extractable fraction of 550 PyOM make dilution a valid possibility in the first few days. To further understand the mechanisms behind the negative priming effect of 550 PyOM on SOC mineralization, additional research is needed. Adsorption isotherms and high-resolution imaging of ^13^C-labeled PyOM surface could help investigate the role of sorption in negative priming. Modifying the surface properties of PyOM and understanding its relation to sorption, as well as experiments with varying addition rates, could also be valuable in determining the relative contributions of dilution vs. sorption.

### 350 PyOM addition increased the abundance of some bacterial taxa

The addition of PyOM had a significant effect on the composition of the bacterial and archaeal community almost immediately (Fig 3.A and 3.C). Given the absence of any pH effects, the shift in communities is most likely primarily a response to the C in PyOM. The shift in communities within the first few days of PyOM addition coincides with a sharp increase in the mineralization of the water-extractable fractions in both 350 and 550 PyOM. As PyOM-C is broken down, it creates a pool of degradation byproducts which can alter the chemical profile of soil C. A significant shift in the C profile over time was observed for 550 PyOM (PERMANOVA, *p_time_* = 0.04) but not for 350 PyOM. We postulate that this shift over time is likely due to microbes preferentially using the easily mineralizable PyOM-C compounds first, leading to an increase in more complex, harder to break down compounds in the soil. Supporting this hypothesis, we identified four features that were specific to the 350 and 550 PyOM-amended soils and never present in any unamended soil samples. These features increased in relative abundance over time in the 350 PyOM-amended soils but declined over time in the 550 PyOM-amended soils (Fig. S5 and Table S3). The chemical formula of these features indicates that they are complex hydrocarbons, which suggests that their relative accumulation in the 350 PyOM-amended soils is due to preferential utilization of easily mineralizable PyOM-C compounds. In the case of 550 PyOM, the limited availability of easily mineralizable C would be more likely to lead to the greater use of these complex C compounds by microbes. Further characterization of these features, including confirming that they are PyOM byproducts and a byproduct of microbial breakdown, is needed to gain deeper insights into microbial utilization of PyOM-C substrates.

In line with these findings, we would expect that microbes using easily mineralizable C would increase in relative abundance first, followed by microbes that have the capacity of using complex aromatic C substrates. We observed an increase in the relative abundance of responsive OTUs belonging to the *Gemmatimonas* and *Pseudonocardia* genera after 18 days of incubation in the 350 PyOM-amended soils (Fig. 5.B), coinciding with a period when the rate of water-extractable PyOM-C mineralization was low (Fig. S6). It is plausible that by this point, the most available C is already mineralized and that these bacteria are benefiting from their ability to utilize the condensed aromatic C in 350 PyOM (23,62). This was also evident in soils amended with 550 PyOM - the only significant increase in relative abundance in 550 PyOM amended soils was observed for an OTU belonging to the genus *Gemmatimonas* on Day 10 (Fig. S7). Notably, the *Pseudonocardia* responsive OTU exhibited 100% BLAST similarity to a *Pseudonocardia* lab isolate that was isolated from burned soils in California on media containing PyOM produced at 350 °C. This provides further evidence for the capacity of *Pseudonocardia* to degrade the carbon present in 350 PyOM.

In contrast, *Noviherbaspirillum* responsive OTUs significantly increased in relative abundance in 350 PyOM amended soils on Day 2 following PyOM addition (Fig. 5.B). Previous incubation studies have observed an increase in the relative abundance of *Noviherbaspirillum* with PyOM, which is attributed to their capacity to degrade aromatic C in PyOM (22,63). Furthermore, *Noviherbaspirillum* species have been found to be more abundant in post-fire soils, suggesting their ability to exploit post-fire resources such as PyOM (64).

### PyOM addition did not result in fungal community shifts

We observed no change in fungal community composition in response to the addition of PyOM (Fig. 3.B and 3.D). This was surprising, given that previous research has shown PyOM to impact whole community composition as well as specific fungal groups (30,65,66). Among the fungal responders, *Calyptrozyma* sp. have been identified as fire-responsive (28,67). Their ability to thrive in post-fire soils is attributed to reduced competition from other fungi and capacity to grow on charred aromatic C. It is possible that bacteria were perhaps better able to utilize the nutrients provided by PyOM, and the effects of PyOM on fungi may only become apparent over longer durations when the easily mineralizable carbon becomes limited. This is supported in many ways by previous studies that have investigated PyOM effects on fungi. For example, Li et al. (32) found that bacteria may be more affected by the aqueous extractable substances of PyOM which could appear over shorter durations, while fungi may be more affected by the porous nature and aromatic carbon compounds. Yu et al. (68) also observed that the PyOM-induced priming effect in their study was strongly associated with the increase of certain bacteria in the first 8 days, with an increase in fungal groups not observed until day 40. Liu et al. (69) found an increase in the proportion of bacteria in fresh PyOM amended soils, and fungi in 6-year-old PyOM amended soils. Another factor that may have contributed to the lack of response in our study is the fine grinding of PyOM particles (chosen in order to ensure effective mixing and even distribution). This may have increased the microporosity, making it more difficult for fungi to colonize the PyOM (32). Additionally, sieving the soil before setting up the incubation could have affected filamentous fungi more than many bacteria (although sieving before incubations is a standard practice to homogenize soil across replicates). More research is needed to fully understand the effects of PyOM on fungal communities, including the role of time and the specific mechanisms at play. Future studies should also consider the potential impacts of particle size and porosity on fungal colonization.

Overall, the effects of PyOM on microbes are likely related to changes in nutrient provision, including both C and N. Other PyOM properties such as surface and electrochemical properties can also affect microbial response to PyOM (70,71). Ash content contributes to the alkalinity of the PyOM and is known to cause small changes in the microbial community composition (59). With our pH adjustment, the effect of ash content should be negligible compared to the effects of PyOM-C. The porous nature of PyOM can adsorb water, organic materials and nutrients, and provide a habitat for microbes (23,72). Furthermore, PyOM sorption of acyl-homoserine lactone (AHL) intercellular signaling molecules can disrupt cell-cell communication among bacteria and affect C mineralization, especially in the short-term following addition of fresh PyOM (21). The porosity and surface area of PyOM increase with pyrolysis temperatures (73), which may influence the mineralization of 350 vs. 550 PyOM. While we did not specifically investigate the microbial colonization of 350 and 550 PyOM or other effects mentioned here, we anticipate that C availability will be the dominant factor controlling microbial response in a 30-day incubation.

## Conclusion

In this study, we demonstrated that the water-extractable carbon in PyOM comprises a small fraction of PyOM-C but exhibits disproportionately high mineralizability. We calculated mineralization rates for the water-extractable PyOM-C fractions of 0.23% day^-1^ for the 350 PyOM and 0.04% day^-1^ for the 550 PyOM. These rates surpass decomposition rates estimated by Wang et al. (7) for wood-based PyOM (mean: 0.004% day^-1^) or PyOM produced within a similar pyrolysis temperature range (mean: < 0.04% day^-1^ for PyOM produced between 200-550 °C), based primarily on lab incubation data. Although estimating these numbers in the field is challenging, some field evidence also exists for a rapidly cycling, easily mineralizable PyOM-C fraction (74,75). Field mineralization rates of PyOM-C may vary, being lower as reported by Maestrini et al. (76), or higher as observed by Major et al. (74) and Keith et al. (12). In the short term, environmental conditions including temperature and precipitation, and soil properties like texture can influence the water-extractable fraction’s fate, determining whether it is transported down the soil profile as leachate, adsorbed to clay minerals, or subject to lateral movement due to erosion. Furthermore, the presence of other labile C sources can also affect PyOM-C mineralization through priming effects (11). Over the long term, ageing-related changes, such as surface oxidation, may make the C in PyOM, particularly in lower temperature PyOM, more susceptible to microbial mineralization (9).

We also observed that the low-temperature 350 PyOM displayed higher C mineralizability than the high-temperature 550 PyOM. Moreover, our short-term incubation study showed net positive priming of SOC upon the addition of 350 PyOM, while the addition of 550 PyOM resulted in net negative priming. Our findings align with (6), indicating that short-term positive priming is primarily attributed to the presence of the water-extractable fraction. This has implications as this fraction can be rapidly transformed and influence soil carbon dynamics, especially in low-C soils that are more vulnerable to positive priming (34). In contrast, high-temperature 550 PyOM exhibited a negative priming effect on SOC mineralization, potentially due to dilution, sorption of SOC on PyOM, or inhibition of microbial activity. In post-burn soils with a higher proportion of low-temperature PyOM, the less aromatic PyOM-C constituents may undergo faster turnover, especially immediately after deposition. This process could lead to a positive priming effect, where the addition of easily mineralizable PyOM-C enhances microbial activity and accelerates short-term SOC mineralization, depending on the fate of the easily mineralizable fraction and other soil and climatic factors. Conversely, in post-burn soils containing a higher proportion of high-temperature PyOM, the slower PyOM-C turnover and negative priming effects may be more likely to contribute to long-term carbon sequestration.

Lastly, our study identified that PyOM addition influenced bacterial community composition, leading to an increased relative abundance of certain bacteria that other studies have suggested as being capable of degrading aromatic C compounds in PyOM. Similar responses have been observed for some of the PyOM-responsive bacteria identified in this study in soils post-burn. Two responsive OTUs belonging to the *Massilia* and *Gemmatimonas* genera had high sequence similarity (> 99% BLAST similarity) to bacteria that were identified as pyrophilous following a prescribed burn at the site where we collected soil samples for this study (77). However, it remains unclear whether these bacteria in our study, and even in the field, increase due to actual consumption of PyOM-C. Techniques like DNA Stable isotope probing (qSIP) (78) with ^13^C-labeled PyOM could help identify the bacteria actively incorporating the C in PyOM, and combining SIP with metagenomics can provide further insights into the functional roles of these incorporators (79).

## Acknowledgements

This work was made possible with the financial support of the Department of Energy through awards DE-SC0016365 and DE-SC0020351. We are thankful to Harry Read for labelling chamber design and construction; Akio Enders for “charcoalator” design and construction; Miranda Sikora and Jamie Woolet for growing the labeled trees; Dana B. Johnson, Mengmeng Luo and Elias Kemna for help with setting up the soil-PyOM incubations; Neem Patel for help with soil collection; Kim Sparks and the Cornell Stable Isotope Laboratory for assistance with PyOM analyses; Wisconsin Department of Natural Resources for providing white pine seedlings; Center for High Throughput Computing (CHTC) for bioinformatics and data analysis (https://chtc.cs.wisc.edu/uw-research-computing/cite-chtc.html)

## Data Availability Statement

The CO_2_ flux and LC-MS data associated with this study are available on ESS-DIVE and can be accessed via https://data.ess-dive.lbl.gov/datasets/doi:10.15485/1985918. The sequencing datasets generated during the current study are available in the NCBI SRA under BioProject ID PRJNA986215 (Submission ID: SUB13560992).

## Author Contributions

The author contributions to the chapter are as follows: study conception and design: NZ, TLW, TDB; data collection: NZ, TDB, MSF; analysis and interpretation of results: NZ, TDB, MSF, MFT, TLW; draft manuscript preparation: NZ; manuscript review and editing: NZ, TDB, MSF, MFT, TLW.

