## Supplemental files for "Soil carbon mineralization and microbial community dynamics in response to PyOM addition"

**Supplemental Note S1.** Hoagland’s solution

Eastern white pine seedlings were grown in a custom ^13^CO2 labelling chamber that was maintained at ~400 ppm CO_2_. During the growing period the seedlings were irrigated alternating between 15-60 mL of distilled H2O and an equivalent amount of a modified Hoagland’s solution. The volume of irrigation received was proportional to the size of the seedling and current moisture levels in the soil. The modified Hoagland’s solution used contained (per L): 2.07 g Ca(NO_3_)_2_*4H_2_O, 1.02 g KNO_3_, 0.49 g MgSO_4_*7H_2_O, 0.23 g NH_4_H_2_PO_4_, 8.94 mg Fe-EDTA, 2.84 mg H_3_BO_3_, 1.8 mg MnCl_2_ *4H_2_O, 0.22 mg ZnSO_4_ * 7H_2_O, 0.1 mg H_2_MoO_4_, and 0.08 mg CuSO_4_ * 5H_2_O.

**Supplemental Note S2.** Additional details on PyOM production

We modified the furnace and adapted the PyOM production design developed by Güereña et al. (1). Briefly, the feedstock was placed in a steel cylinder inside the furnace chamber and subjected to a continuous argon gas supply at a rate of 1 L min^‑1^ to maintain anaerobic conditions during pyrolysis. The heating rate for production of PyOM was kept constant at 5 °C min^−1^ and mixing of the feedstock inside the steel cylinder via mechanical paddles started once temperatures hit 250 °C. We held the temperature constant for 30 min once the target temperature was reached, after which the PyOM was rapidly cooled by circulating cold water in stainless steel tubes wrapped around the steel cylinder.

**Supplemental Note S3.** Additional details on water-extractable PyOM-C extraction and exchange

To separate the water-extractable PyOM-C, we used a series of five sequential extractions with ultrapure water, using the methods described in Smith et al. (2) and Whitman et al. (3). In each round of extraction, PyOM was shaken with ultrapure water (1:10 solid:water ratio) in a 15 mL Falcon conical centrifuge tube (Corning Inc., Corning, NY, United States) for 15 min and then centrifuged at 2593 x *g* for 15 min at 4 °C. After centrifugation, the supernatant was syringe-filtered through a sterile 0.45 μm C-free glass microfiber filter (cat # 6902-2504, Cytiva, Palo Alto, CA, United States). The resulting extracts were retained, with sequential extractions performed on the remaining PyOM and pooled in order to extract as much DOC as possible. The total dissolved organic carbon (DOC) content of the water-extractable PyOM-C fraction was measured at the Water Science and Engineering Laboratory, UW-Madison using a Sievers M5310C Total Organic Carbon Analyzer (SUEZ Water Technologies & Solutions, Trevose, PA, United States).

**Supplemental Note S4.** Additional details on ^13^CO_2_ partitioning

We determined the fraction of CO_2_ emitted from non-water-extractable PyOM (*f_non-water-extractable PyOM_*) by partitioning the total CO_2_ emissions from the “Soil + ^13^C non-water-extractable PyOM” treatments into two sources: CO_2_ emitted from the ^13^C labeled non-water-extractable PyOM source and CO2 emitted from the unlabeled soil and water-extractable PyOM source, as represented by Equation 1:

(1) *f_non-water-extractable PyOM_ = (δ_Total CO2_ – δ_CO2 soil+water-extractable PyOM_)/ (δ_CO2 non-water-extractable PyOM_ - δ_CO2 soil+water-extractable PyOM_)*

In this equation, *δ_Total CO2_* represents the mean isotopic composition of the total CO_2_ emitted from the "Soil + ^13^C non-water-extractable PyOM" treatments, *δ_CO2 soil+water-extractable PyOM_* represents the mean isotopic composition of the CO_2_ emitted from soil and water-extractable PyOM fraction, determined from "Soil + unlabeled PyOM" treatments, and *δ_CO2 non-water-extractable PyOM_* represents the isotopic composition of the CO_2_ emitted from non-water-extractable PyOM, which was assumed to be the isotopic composition of the ^13^C labeled PyOM.

We determined the fraction of CO_2_ emitted from SOC (*f_SOC_*) by partitioning the total CO_2_ emissions from the “Soil + ^13^C PyOM” treatments into two sources: CO_2_ emitted from the ^13^C labeled PyOM source and CO_2_ emitted from the unlabeled soil source, as represented by Equation 2:

(2) *f_SOC_ = (δ_total CO2_ – δ_CO2 PyOM_)/ (δ_CO2 SOC_ - δ_CO2 PyOM_)*

In this equation, *δ_total_* represents the isotopic composition of the total CO_2_ emitted from "Soil + ^13^C PyOM" treatments, *δ_CO2 PyOM_* represents the isotopic composition of the CO_2_ emitted from PyOM, which was assumed to be the isotopic composition of the ^13^C labeled PyOM, and *δ_CO2 SOC_* represents the isotopic composition of the CO_2_ emitted from SOC, determined from the "Unamended soil control" treatments.

| **Table S1.** Properties of biomass | | | | |
| --- | --- | --- | --- | --- |
| Property (units) | ^13^C labeled biomass | | Unlabeled biomass | |
|  | Stems | Needles | Stems | Needles |
| Total C (%) | 41.56 ± 0.92 | 41.61 ± 2.05 | 47 ± 0.46 | 47.02 ± 1.05 |
| Total N (%) | 1.50 ± 0.03 | 2.34 ± 0.11 | NA | NA |
| Bulk δ^13^C vs. VPDB (‰) | 914.93 ± 3.62 | 2071.03 ± 9.69 | -28.65 ± 0.29 | -29.96 ± 0.32 |
| Bulk δ^15^N vs. atmospheric air (‰) | 10.17 ± 4.64 | -2.16 ± 0.30 | NA | NA |
| The values presented are means of five replicates ± standard deviation. | | | | |

| **Table S2.** Properties of non-water-extractable PyOM | | | | |
| --- | --- | --- | --- | --- |
| Property (units) | ^13^C labeled | | Unlabeled | |
|  | 350 °C | 550 °C | 350 °C | 550 °C |
| pH (H_2_O) | 7.96 | 9.24 | 7.02 | 9.95 |
| Total C (%) | 64.71 ± 1.06 | 75.17 ± 1.9 | 68.64 ± 2.65 | 78.18 ± 0.52 |
| Total N (%) | 3.28 ± 0.04 | 3.05 ± 0.05 | 2.69 ± 0.09 | 2.50 ± 0.02 |
| Bulk δ^13^C vs. VPDB (‰) | 1518.48 ± 2.34 | 1616.97 ± 7.39 | -29.14 ± 0.14 | -29.41 ± 0.02 |
| Bulk δ^15^N vs. atmospheric air (‰) | 0.25 ± 0.29 | 0.39 ± 0.43 | 0.16 ± 0.15 | 0.72 ± 0.29 |
| Total organic C (%) | 56.56 ± 0.23 | 62.62 ± 0.2 | 59.25 ± 0.31 | 65.13 ± 0.61 |
| pH values of the original PyOM before adjustment are shown. The values presented for total C, N, δ^13^C and δ^15^N are means of five replicates ± standard deviation. The values for total organic C represent means of three replicates ± standard deviation. | | | | |

| **Table S3.** Properties of LC-MS features identified in PyOM amended soils | | | |
| --- | --- | --- | --- |
| Feature | Average Rt (min) | Average Mz | Formula |
| 1 | 2.609 | 409.16467 | C_25_H_20_N_4_O_2_ |
| 2 | 2.609 | 425.13583 | NULL |
| 3 | 2.607 | 432.23819 | C_22_H_30_O_6_ |
| 4 | 2.609 | 387.18195 | C_23_H_22_N_4_O_2_ |

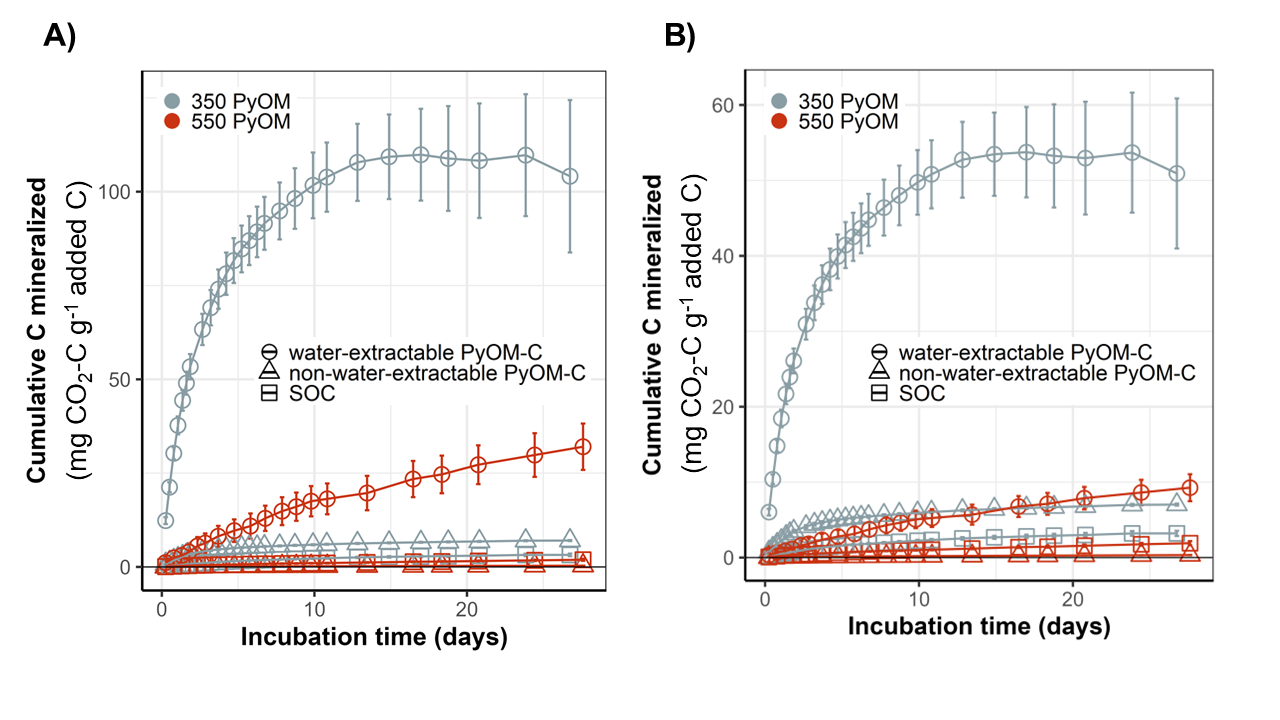

***Figure S1. Mineralizability of different fractions of PyOM over*** ***time when*** ***the labeled water-extractable PyOM-C fraction is*** ***A)*** *depleted in ^13^C than bulk PyOM by 35% and* ***B)*** *enriched in ^13^C than bulk PyOM by 35%. Data in both panels show cumulative mean C mineralized per gram of added C for water-extractable PyOM-C, non-water-extractable PyOM-C and SOC. (n=4-5, error bars=SE)*

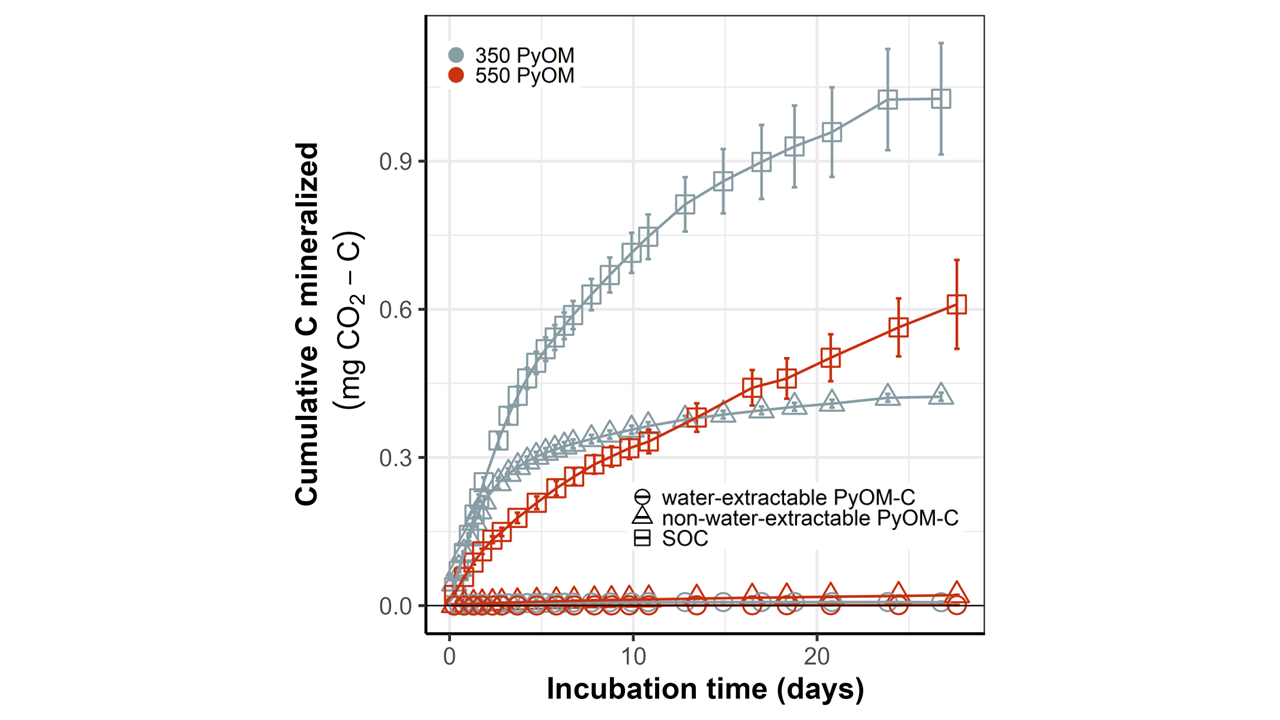

***Figure S2. Mineralization of different fractions of PyOM over time.*** *Cumulative mean C mineralized for water-extractable PyOM-C, non-water-extractable PyOM-C and SOC. (n=4-5, error bars=SE).*

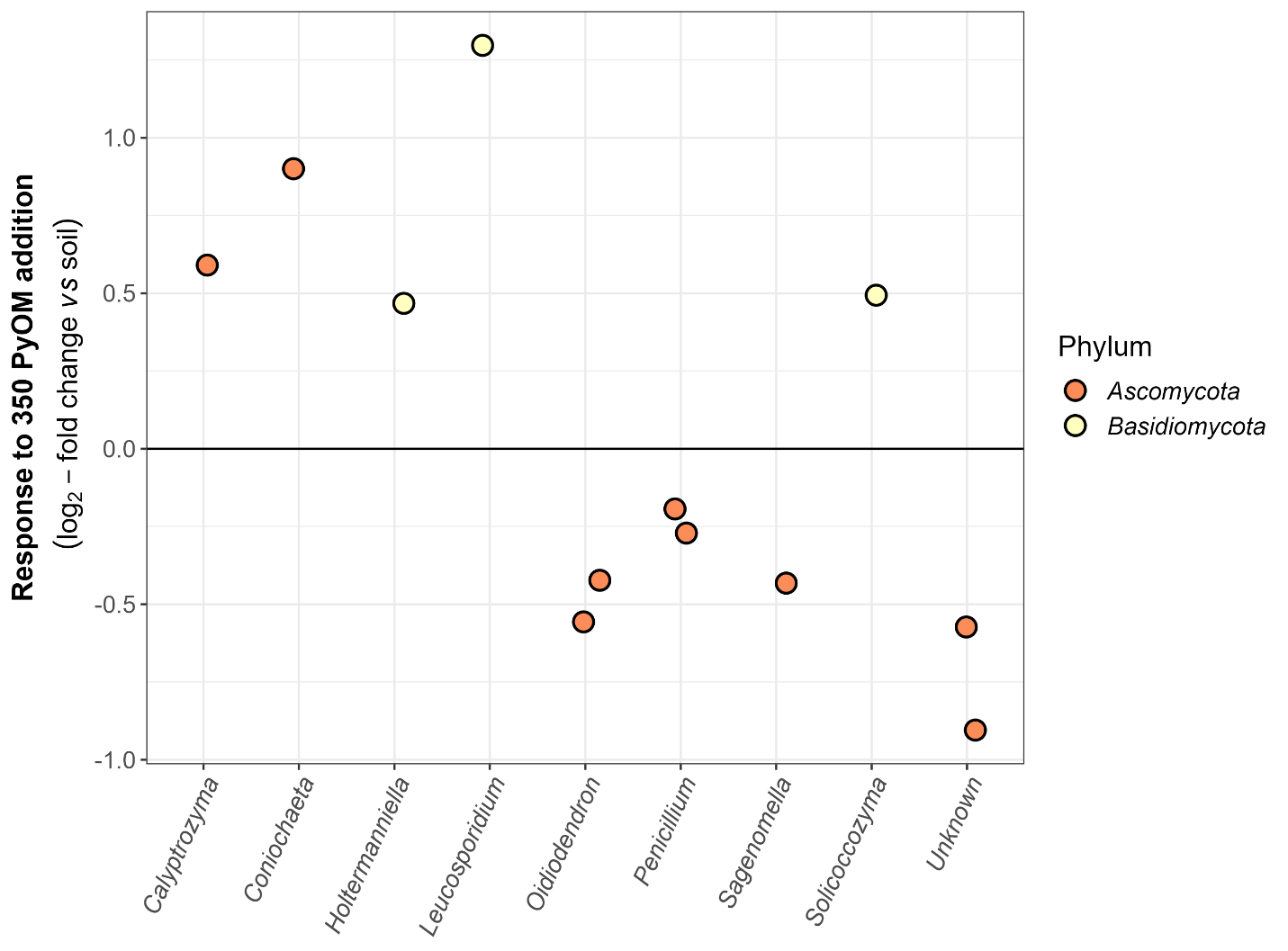

***Figure S3. Fungal response to 350 PyOM.*** *Log2-fold change in 350 PyOM amended vs. unamended soils, controlling for* *differences in taxon abundance across samples on Day 0 and over time.* *Each point represents a single ITS2 gene OTU with mean normalized count above the 25^th^ percentile and that was significantly different in abundance in PyOM amended vs. unamended soils (Benjamini and Hochberg correction, adjusted p value < 0.05).*

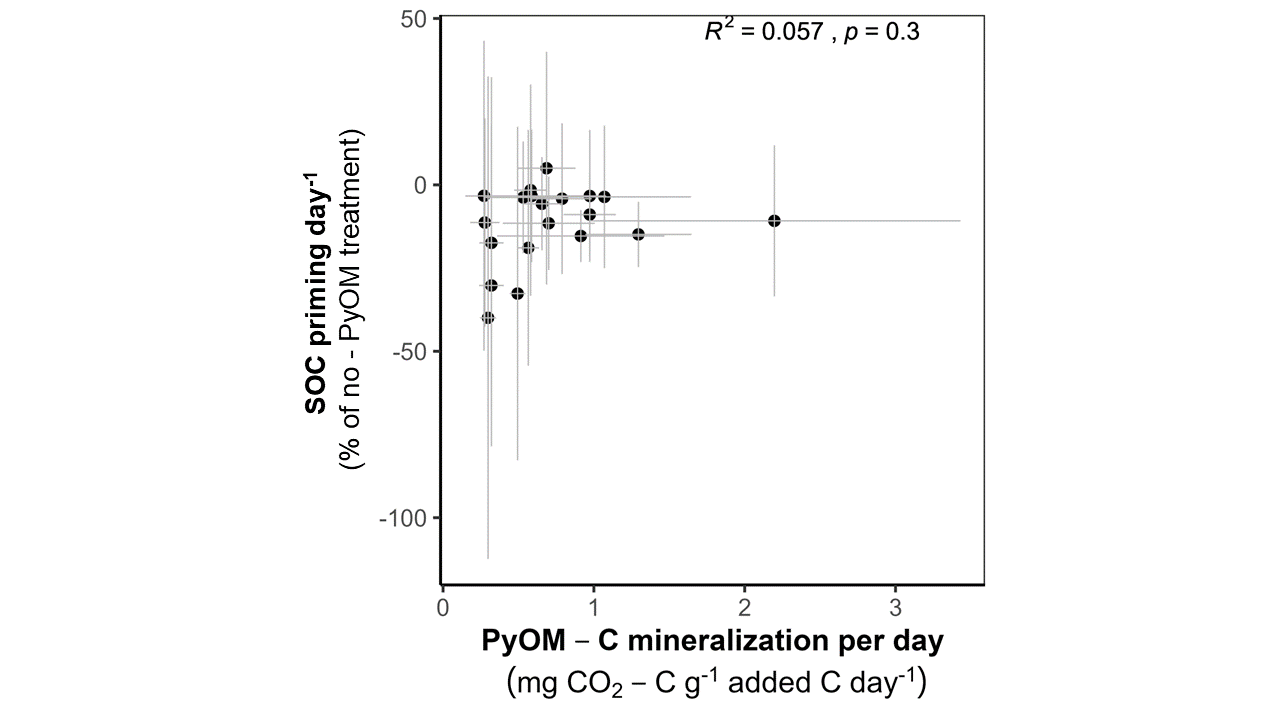

***Figure S4.*** *Relationship between mean PyOM-C mineralization rate and SOC priming for 550 °C PyOM. (n=4-5, error bars=SE)*

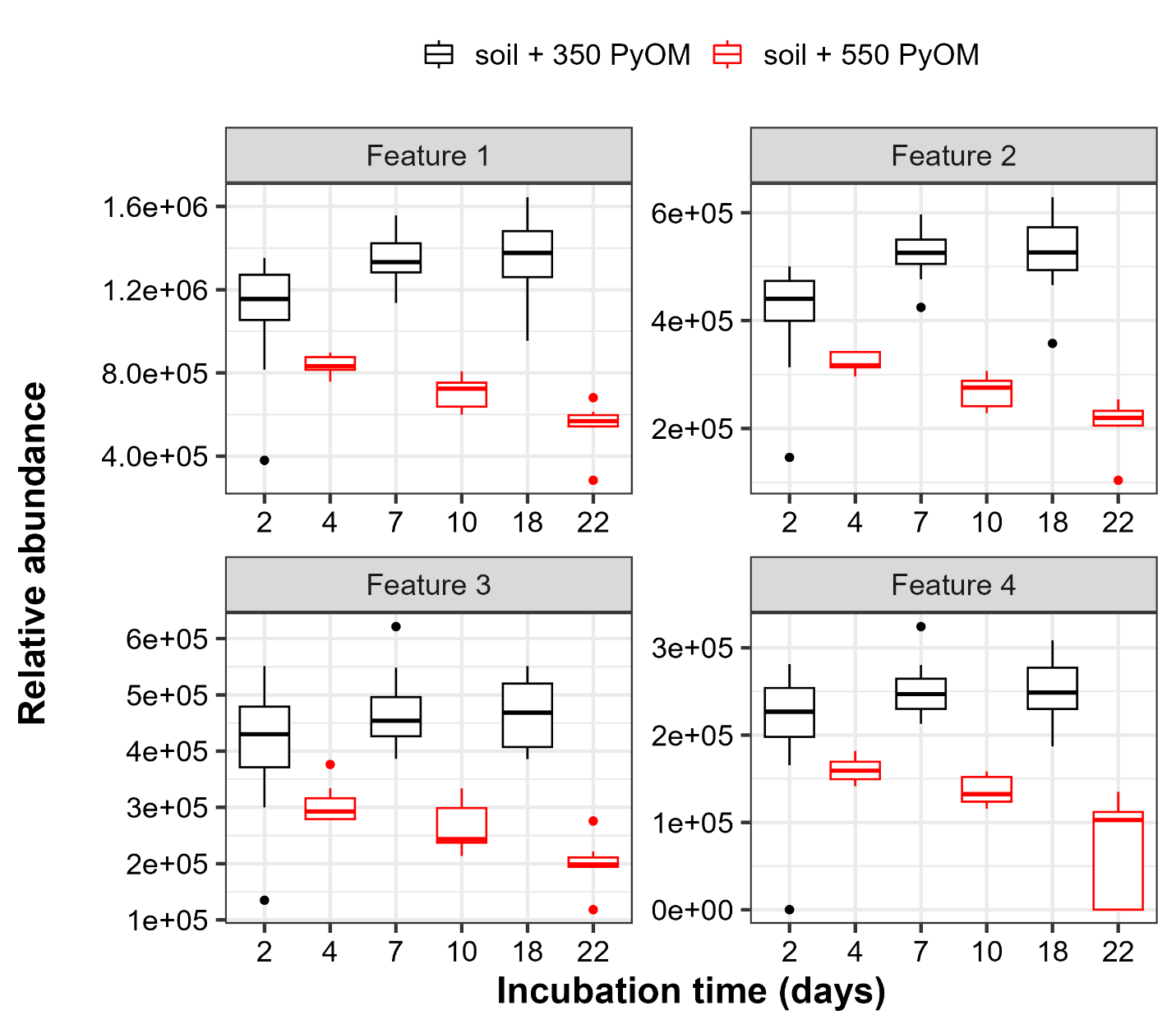

***Figure S5****.* ***LC-MS features identified in PyOM amended soils.*** *Relative abundance of four LC-MS peaks over time observed only in the PyOM-amended soils (n=5-8). Colors indicate soils amended with 350 PyOM (black) and 550 PyOM (red).*

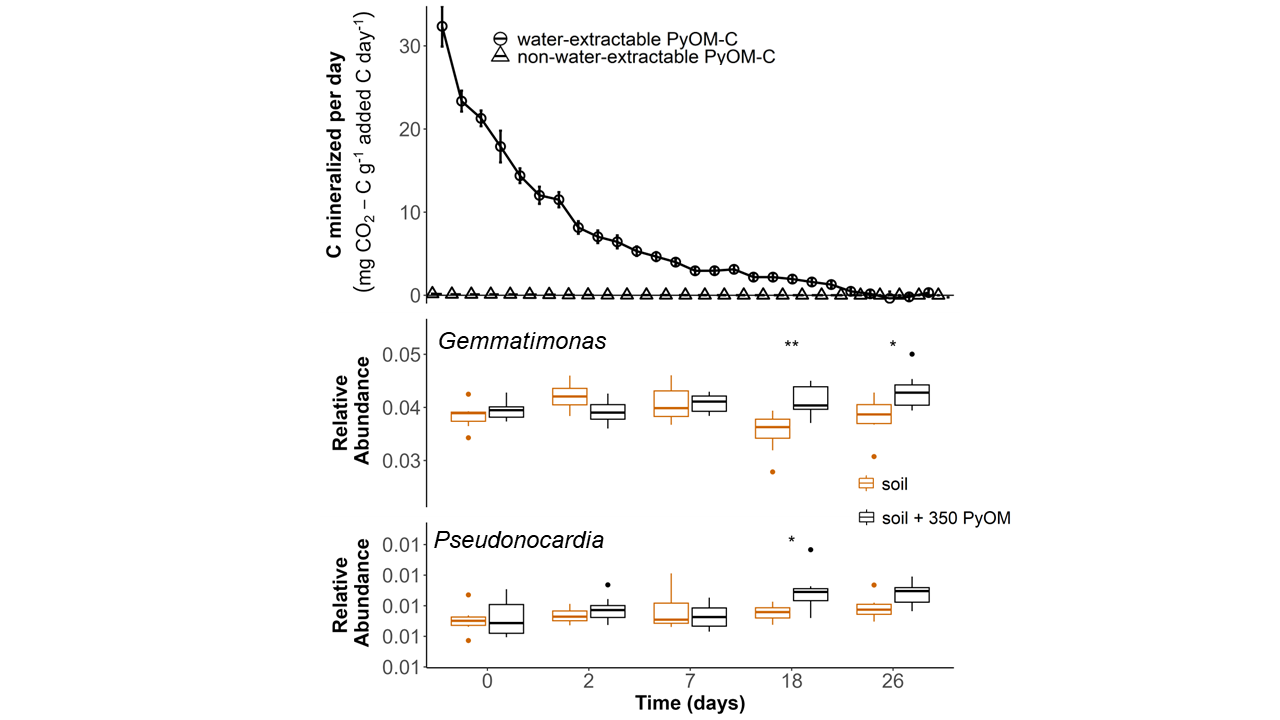

***Figure S6. Late responders to 350 PyOM. A)*** *Rate of 350 PyOM-C mineralization for water-extractable and non-water-extractable fractions (n=4-5, error bars=SE).* ***(B & C)*** *Relative abundance of positive responsive genera* Gemmatimonas *and* Pseudonocardia *over time observed in the unamended and 350 PyOM-amended soils (n=5-8). * indicates relative abundances that differ significantly from unamended soil at a given timepoint (t-test, p < 0.05).*

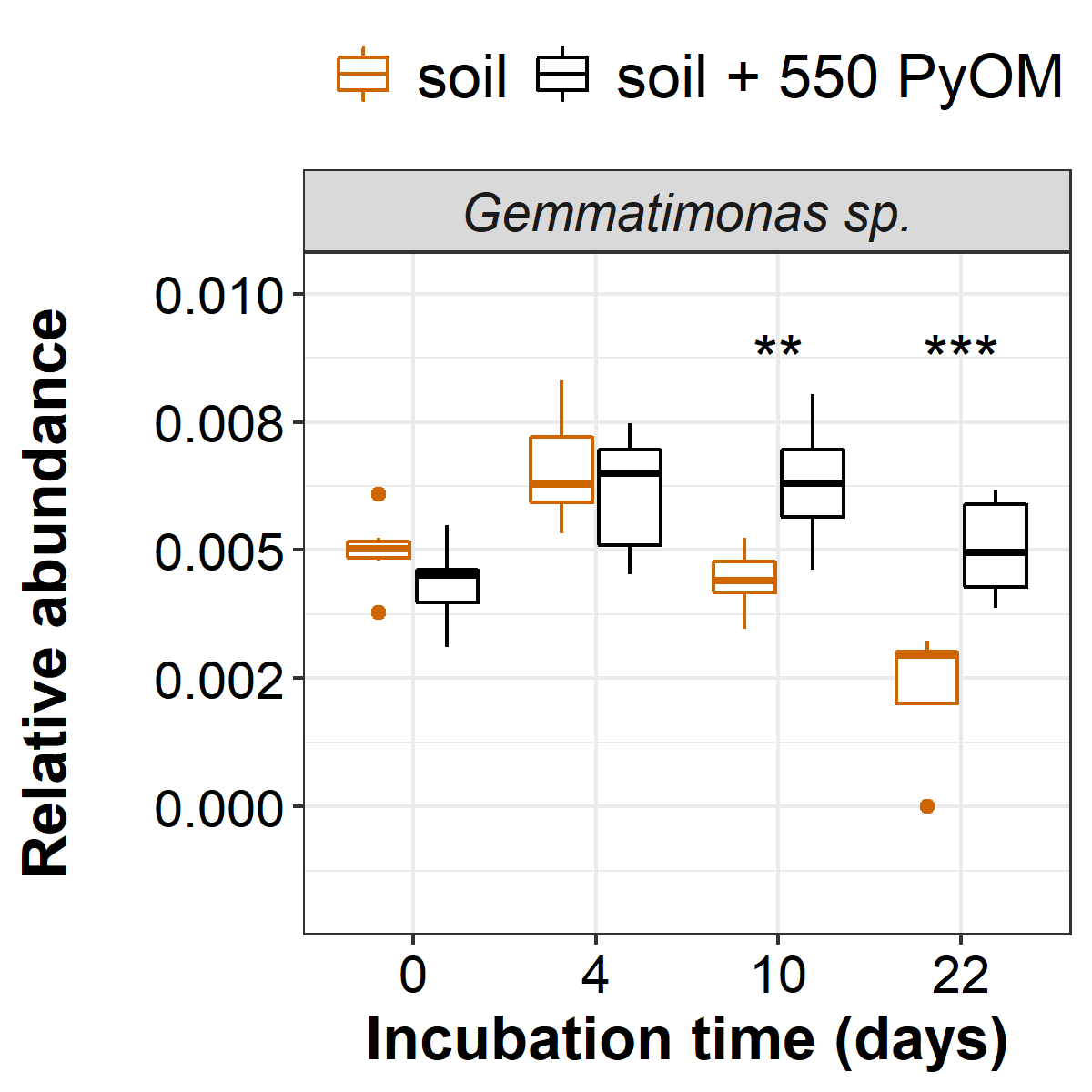

***Figure S7. Response of* Gemmatimonas sp. *to 550 PyOM.*** *Relative abundance of OTU belonging to genus* Gemmatimonas *over time in the unamended, and 550 PyOM-amended soils (n=5-8). * indicates relative abundances that differ significantly from unamended soil (t-test, *: p < 0.05; **: p < 0.01; ***: p < 0.001).*
